# The latitudinal diversity gradient in brush-footed butterflies (Nymphalidae): conserved ancestral tropical niche but different continental histories

**DOI:** 10.1101/2020.04.16.045575

**Authors:** Nicolas Chazot, Fabien L. Condamine, Gytis Dudas, Carlos Peña, Pavel Matos-Maraví, Andre V. L. Freitas, Keith R. Willmott, Marianne Elias, Andrew Warren, Kwaku Aduse-Poku, David J. Lohman, Carla M. Penz, Phil DeVries, Ullasa Kodandaramaiah, Zdenek F. Fric, Soren Nylin, Chris Müller, Christopher Wheat, Akito Y. Kawahara, Karina L. Silva-Brandão, Gerardo Lamas, Anna Zubek, Elena Ortiz-Acevedo, Roger Vila, Richard I Vane-Wright, Sean P. Mullen, Chris D. Jiggins, Irena Slamova, Niklas Wahlberg

## Abstract

The latitudinal diversity gradient (LDG) is arguably one of the most striking patterns in nature. The global increase in species richness toward the tropics across continents and taxonomic groups stimulated the formulation of many hypotheses to explain the underlying mechanisms of this pattern. We evaluated several of these hypotheses to explain spatial diversity patterns in the butterfly family, Nymphalidae, by assessing the contributions of speciation, extinction, and dispersal to the LDG, and also the extent to which these processes differ among regions at the same latitude. We generated a new, time-calibrated phylogeny of Nymphalidae based on 10 gene fragments and containing *ca*. 2,800 species (∼45% of extant diversity). Neither speciation nor extinction rate variations consistently explain the LDG among regions because temporal diversification dynamics differ greatly across longitude. For example, we found that Neotropical nymphalid diversity results from low extinction rates, not high speciation rates, and that biotic interchanges with other regions were rare. Southeast Asia was also characterized by a low speciation rate but, unlike the Neotropics, was the main source of dispersal events through time. Our results suggest that global climate change throughout the Cenozoic, particularly during the Eocene-Oligocene transition, combined with the conserved ancestral tropical niches, played a major role in generating the modern LDG of butterflies.

## INTRODUCTION

Understanding the uneven distribution of biodiversity on Earth is one of the most fundamental goals in ecology and evolution. Numerous patterns of biodiversity distributions have been documented, but the obvious increase in species richness from the poles towards the equator known as the latitudinal diversity gradient (LDG) is remarkable for its consistency across geographic scales and taxonomic groups (Mittelbach et al. 2007, Mannion et al. 2014, Kinlock et al. 2017). Although many different hypotheses have been formulated to explain this pattern, no consensus has emerged.

The proposed hypotheses fall into three broad categories: ecological, evolutionary, and historical (Mittelbach et al. 2007). The increasing availability of molecular phylogenies has renewed interest in evolutionary and historical hypotheses because they provide an opportunity to infer some of the past history without extensive fossil information (Wiens et al. 2006, Wiens et al. 2009, Jansson et al. 2013, Economo et al. 2018). Four historical processes that could result in greater species richness in tropical regions are usually proposed:

1. Longer time-for-speciation in the tropics (Willis 1922, Stephen & Wiens 2002, Wiens & Donoghue 2004). During the early Cenozoic, tropical biomes were found across much higher latitudes, while colder biomes with higher seasonality extended only after the Eocene-Oligocene boundary. Many groups from the early Cenozoic probably originated in these tropical areas. Assuming similar speciation and extinction rates across regions, species richness would therefore be greater in the tropics if lineages had more time to accumulate.
2. Asymmetric dispersal events between the tropics and other areas. Clades originated either in the tropics and rarely dispersed out of them (Farrell 1992, Jablonsky et al. 2006), or in temperate regions and frequently dispersed into the tropics, thereby increasing tropical species richness (*e.g*., Condamine et al. 2012). The first scenario is expected in the cases where most tropical organisms highly specialised to their environmental niche cannot colonize different ecological conditions such as those in seasonal temperate regions. Consequently, such colonization events may be rare and recent, resulting in strong conservatism of the tropical niche. The second scenario implies that clades originated in temperate regions but colonized tropical regions frequently, resulting in repeated evolution of adaptations for tropical existence.
3. Higher speciation rates in the tropics (“cradle of diversity”, Fisher 1960). Tropical lineages speciate more rapidly than temperate lineages. Proposed mechanisms that promote high speciation rates in tropical regions include: larger area (a species-area effect; Rosenzweig 1995), faster evolutionary rate (through the effect of temperature on mutation rate and generation times) (Allen et al. 2006), and increased biotic interactions (Fischer 1960, Schemske 2002).
4. Lower extinction rates in the tropics (“museum of diversity”, Fisher 1960, Rosenzweig 1995). Tropical regions are perceived as more stable and less prone to drastic climate change (*e.g*., Janzen 1967), thereby reducing extinction risk. Further, it has been argued that larger species ranges in tropical areas permit larger population sizes, which reduce species extinction risk (Fisher 1960).

Tests of these hypotheses have focused primarily on vertebrates and plants, and with a few exceptions (*e.g*., Condamine et al. 2012, Economo et al. 2018) large, densely sampled phylogenetic trees with robust divergence time estimates have been lacking for insects, the most species-rich terrestrial animal group. Here, we generated the first species-level phylogeny of the brush-footed butterflies (Nymphalidae), the most diverse butterfly family (∼6400 described species). Over the past two decades, a sustained effort has been made to generate comparable molecular data across the family, such that we can now assemble a densely sampled phylogenetic tree. We aggregated data from virtually all nymphalid species ever sequenced to date to generate a time-calibrated tree of 2800 species, representing about 45% of the extant described species. Nymphalid butterflies appear to have originated in the Late Cretaceous (Wahlberg et al. 2009, Chazot et al. 2019) and diversified across all continents. The family exemplifies a latitudinal diversity gradient with ∼83% of described species distributed in the Neotropics, Afrotropics, or Southeast Asian biogeographic regions, while the Palearctic and Nearctic regions together account for ∼15% of the total species richness. Here, we assess the relative contribution of each of the four proposed mechanisms (longer time-for-speciation, dispersal variation, higher tropical speciation rates, and lower tropical extinction rates) in generating the modern LDG of nymphalids using the new time-calibrated phylogeny and biogeographic information. These four mechanisms however, usually assume a binary model in which processes occurring at the same latitude are homogenous but differ from processes occurring at different latitudes. Yet, the dynamics of diversification and the underlying processes may differ widely among regions for historical reasons (*e.g*., different colonization times) or geologic and climatic features (*e.g*., Andean uplift in South America). Accordingly, we investigated the extent to which age, dispersal, speciation, and extinction differ longitudinally across the tropical regions.

## RESULTS

### Time-calibrated supertree

Nymphalidae diverged from its sister clade (Riodinidae + Lycaenidae) 93.2 [84.4 – 101.8] Myr ago, with a crown age of 84.6 [76.0 – 91.8] Myr ago for nymphalids (Figure 1, Supplementary Information I & II). This age is within the range of previously inferred ages for the family. Our source of secondary calibrations (Chazot et al. 2019) found a crown age of 82.0 [68.1 – 98.3] Myr ago, an estimate similar to Wahlberg et al. (2013), Heikkilä et al. (2011), Espeland et al. (2018) and Condamine et al. (2018, one clock). Note that Wahlberg et al. (2009) found a mean crown age about 12 million years older. The backbone topology of our tree agreed with previous studies, but the position of Libytheinae was poorly supported. The taxon is often recovered as sister to all other Nymphalidae. A study with substantially more genetic loci (352 markers) did not increase support for its position within the family (Espeland et al. 2018).

**Figure 1.**
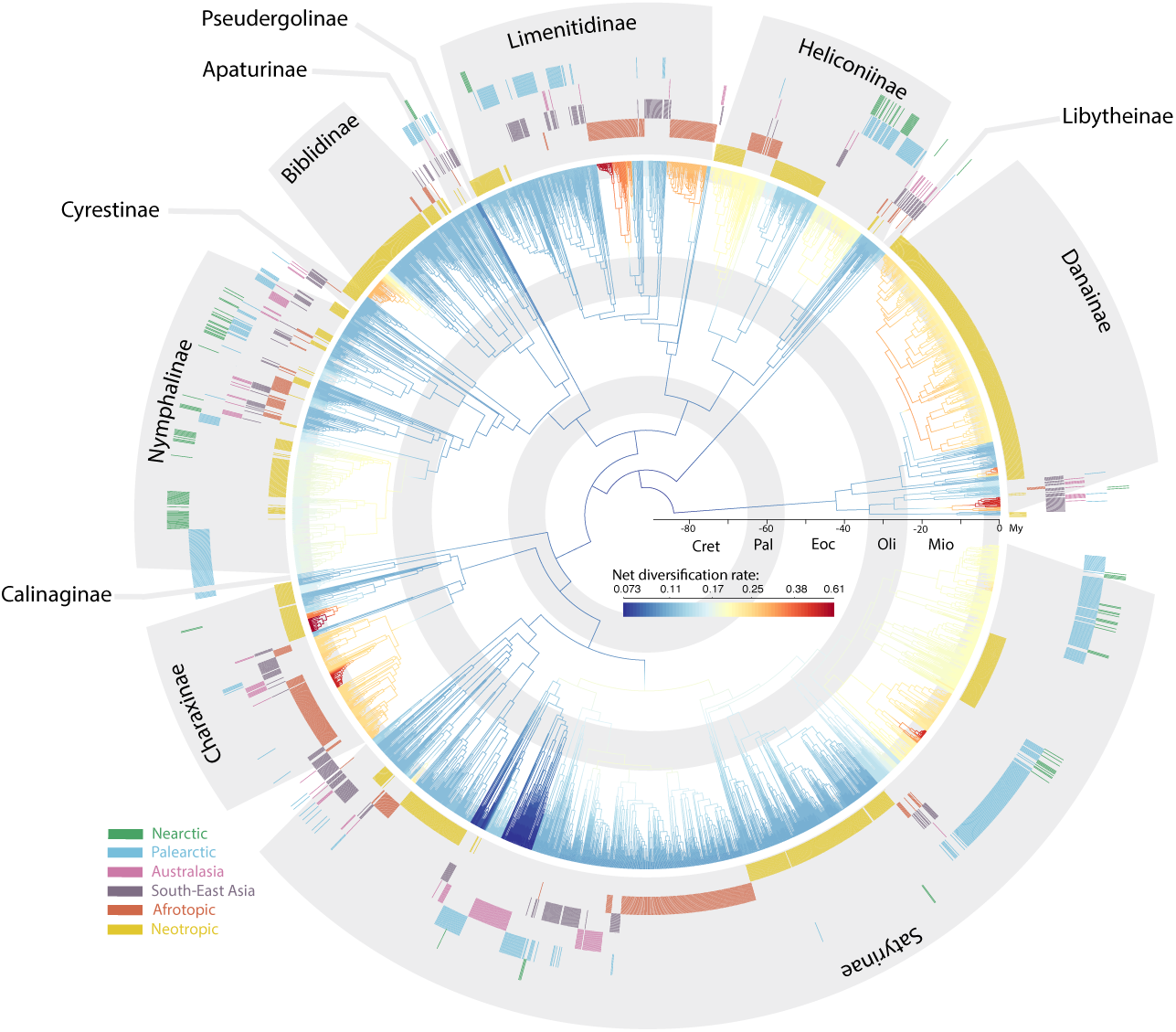
Time-calibrated phylogenetic tree of brush-footed butterflies (Nymphalidae) and biogeographic distribution of extant species (coloured bars in outer circles indicate the biogeographic distribution of each terminal). Branches are coloured according to the average posterior net diversification rate from a birth-death analysis of BAMM. Subfamilies are indicated outside the tree. Grey circles inside the phylogeny indicate geological time periods: Cret = Cretaceous, Pal = Paleocene, Eoc = Eocene, Olig = Oligocene, Mio = Miocene.

### Biogeographic patterns of diversification: a whole-tree approach

Ancestral area estimations with a maximum-likelihood Dispersal-Extinction-Cladogenesis (DEC, Ree & Smith 2008) model inferred an ancestral range at the root of Nymphalidae covering Southeast Asia, Palearctic and western Nearctic in the Cretaceous (Supplementary Information IV, V, VI). We remain cautious about this result because long branches associated with widespread groups such as the Libytheinae and Danainae can be problematic for ancestral range estimations (Crisp et al. 2005). Nevertheless, early lineages diversified almost entirely in Southeast Asia before they dispersed towards the Afrotropics and Neotropics by the end of the Paleocene (66-56 Myr ago, Supplementary Information VI). During the Eocene (56-34 Myr ago), Southeast Asia became even more central to the dispersal of Nymphalidae with nearly 60% of intercontinental dispersal events originating from that region (Supplementary Information V). Many dispersal events dated to the Eocene occurred between low latitude tropical regions. During the Oligocene (34-23 Myr ago), Southeast Asia remained the most common origin of dispersal events between regions with almost 30% (Supplementary Information V). However, during this epoch we inferred more frequent dispersal events from the Afrotropics into Southeast Asia (*ca*. 10%). Compared to earlier periods, the Neotropics became increasingly isolated, whereas interchanges continued between Southeast Asia, Australasia, the Afrotropics, and the Palearctic, a pattern that was strengthened during the Miocene (Supplementary Information V). Three types of dispersal events prevailed in the Miocene: from Southeast Asia towards Australasia (*ca*. 24% of the dispersal events—twice as frequent as during the Oligocene), from Southeast Asia towards the Palearctic (*ca*. 18%), and from the Palearctic towards the Nearctic (*ca*. 17%).

The average net diversification rate (speciation minus extinction) across nymphalid butterflies increased through time globally and in all regions except Australasia (Figures 1 & 2, Supplementary Information III, IV, V, VI). We also found evidence for contrasting patterns of speciation and extinction rates through time among different tropical regions. Compared with other regions, the average Neotropical speciation rate was high during the Eocene, but not during the last 40 Myr (Figure 2A). In parallel, we found the lowest average extinction rate in the Neotropics during the Eocene, Oligocene, and Miocene. By contrast, average speciation and extinction rates increased rapidly in the Afrotropics, especially during the Miocene (Figure 2B, C), but the overall net diversification rate increased through time. On average, Southeast Asian lineages diversified more slowly than other tropical regions (Figure 2 & 3), but Australasia showed a distinctly different pattern (Figure 2). Australasia was the only region characterized by decreasing speciation and net diversification rates. Both were particularly high during the Eocene, but decreased rapidly at the end of the Eocene and continued until the Pliocene. The Palearctic region was characterized by a rapid increase in speciation rate and net diversification at the end of the Eocene, the fastest diversification rate of any region during the Oligocene. Speciation and net diversification diminished during a period corresponding to the mid-Miocene climatic optimum (*ca*.15 Myr ago) before increasing again. The Nearctic region, which is the least diverse region despite having been colonized early in the history of Nymphalidae, hardly deviates from steady increases in speciation and net diversification rates. However, the Oligocene extinction rate was among the highest.

**Figure 2.**
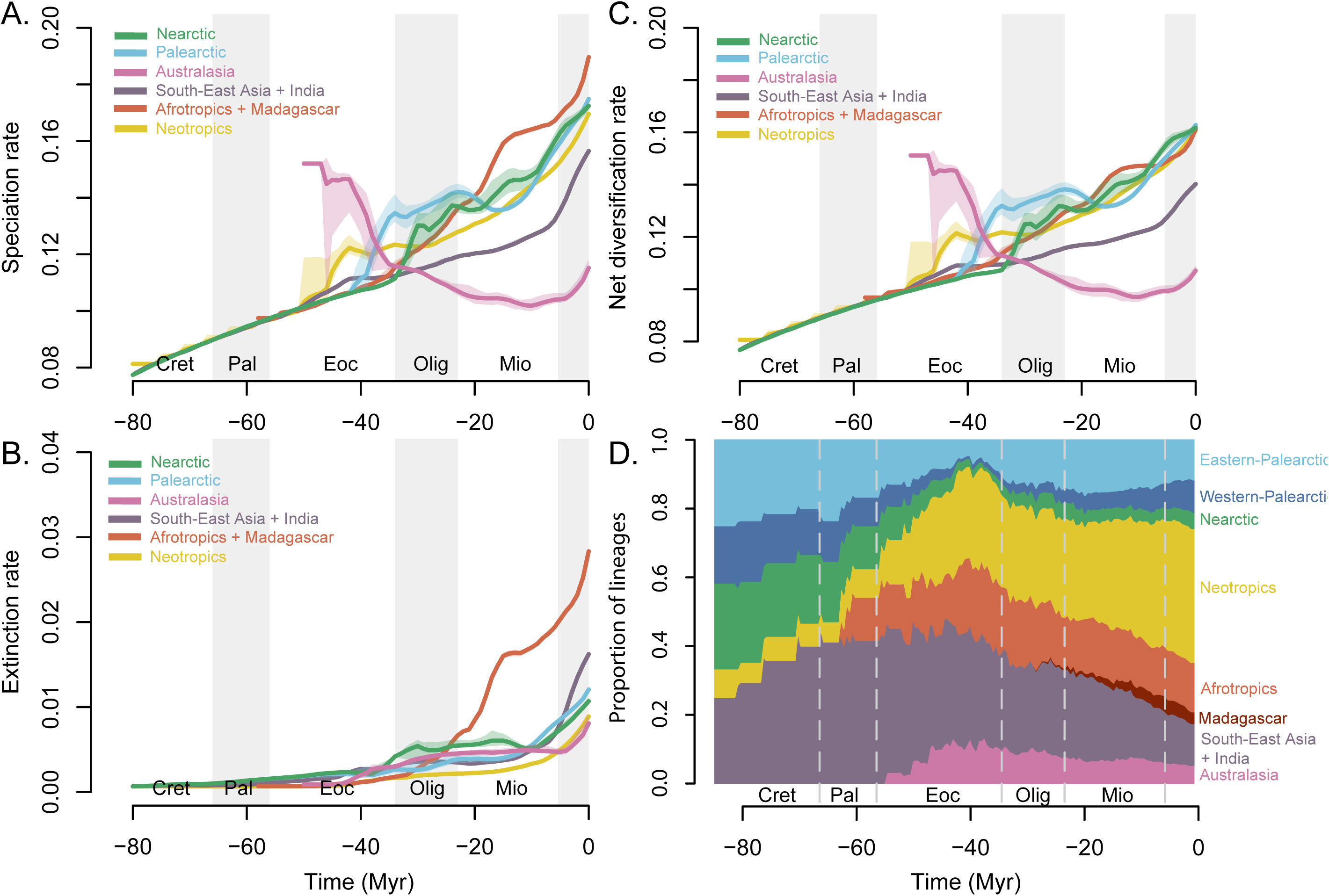
Temporal dynamics of speciation (A.), extinction (B.), and net diversification (C.) rates in each region. Rates were estimated using a sliding window analysis combining historical biogeography (DEC) and speciation/extinction rates (BAMM). Coloured shading indicates the distribution of mean rates estimated for 100 randomly sampled timing of dispersal events. Coloured lines are the mean of this distribution. D. Relative proportion of lineages in each biogeographic region through time, estimated using our DEC results. Cret = Cretaceous, Pal = Paleocene, Eoc = Eocene, Olig = Oligocene, Mio = Miocene.

**Figure 3.**
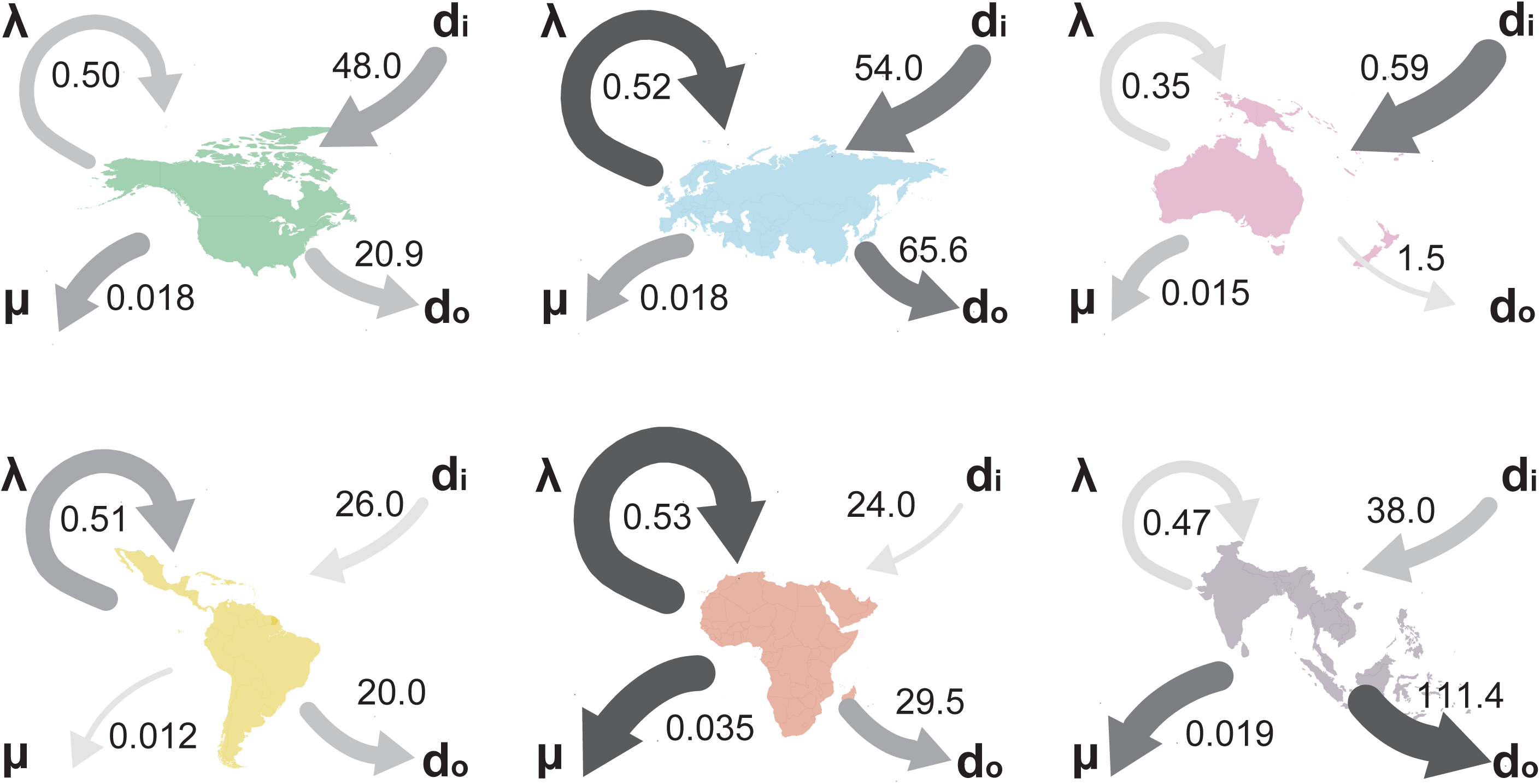
Relative contributions of speciation rate (λ), extinction rate (μ), dispersal into (d_i_), and dispersal out (d_o_) across six biogeographic regions. Colour and thickness of the arrows are proportional to their relative values. For instance, Southeast Asia (Sa) has the highest d_o_ of all regions but among the lowest λ. Each of these statistics where estimated within four large time intervals (Cretaceous-Paleocene; Eocene; Oligocene; Miocene-Present) and summed.

Comparing the relative frequency of lineages sampled in the tree in each biogeographic region, we found the Eocene to be a period of major transition. Neotropical, Afrotropical, and Australasian lineages increased in relative proportion to Nearctic and Palearctic lineages (Figure 2D). Strikingly, Nearctic and Palearctic lineages reached their lowest historical relative frequency at the end of the Eocene before starting to diversify to their modern extent.

### Biogeographic patterns of regional diversification

We compared the diversification dynamics among continents, and focused on clades including at least four sampled lineages having mostly diversified in a single region (hereafter called regional diversification events, Supplementary Information VII). We identified 90 regional diversification events. Thirty of these clades are Southeast Asian or Australasian, 21 are Afrotropical, 21 are Neotropical, 14 are Palearctic, and four are Nearctic. We extracted the crown age and inferred net diversification at the crown (netDiv_crown_) and net diversification at present (netDiv_0_) from time-dependent birth-death diversification models fitted to each clade. Nearctic diversification events were clearly the youngest (Figure 4A; Supplementary Information VII). Southeast Asian + Australasian nymphalid clades showed the widest range of crown ages, ranging from 1.37 – 43.87 Ma (Figure 4A.). The Neotropics were also characterized by a wide range of crown ages but, most importantly, by the oldest radiations on average. The Afrotropics were characterized by the widest range of net diversification parameters at present (netDiv_0_), while Southeast Asia + Australasia was characterized by the widest range of net diversification rates parameter at the origin of the clade (netDiv_crown_). Coefficients of time variation (noted α), show a strong pattern of decreasing speciation rate through time among the four Nearctic diversification events and to a lesser extent the Palearctic clades (Supplementary Information VII). The other three regions showed relatively balanced distributions of α, demonstrating no consistent change in speciation rate through time.

**Figure 4.**
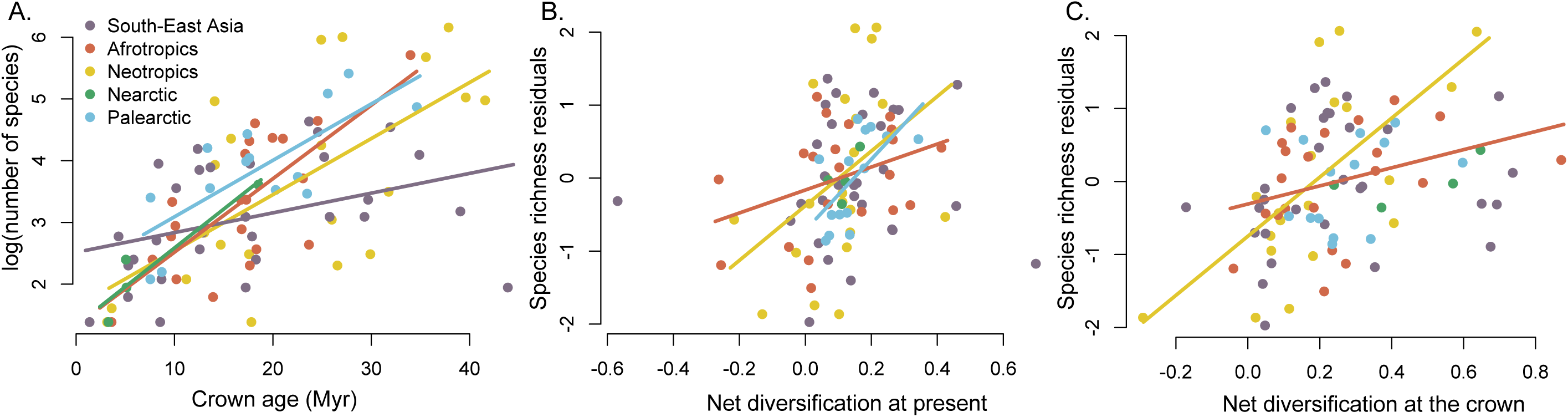
Analyses of 90 regional diversification events. A. Number of species (log scale) in each diversification event as a function of its crown age (Myr). B. Residual species richness after removing the effect of time as a function of the net diversification rate at present. Only relationships within the Palearctic, Afrotropical, and Neotropical regions were significant. C. Residual species richness after removing the effect of time as a function of the net diversification at the crown. Only relationships in the Afrotropics and the Neotropics were significant. Regression lines with significant slopes are shown.

We found a strong positive relationship between the crown age of regional diversification events and extant species richness, except in the Nearctic (Figure 4A.). NetDiv_0_ and netDiv_crown_ also had significant effects on species richness in the Neotropics, Afrotropics, and the Palearctic (Supplementary Information VII). We repeated these tests on the residuals of a linear regression between the crown ages and species richness of clades to determine whether differential speciation or extinction explained region-specific patterns after removing the effect of time. In the Neotropics and Afrotropics, both netDiv_0_ and netDiv_crown_ had a significant effect on explaining residual species richness (Figure 4B, C). In the Palearctic, the recent dynamics of diversification (netDiv_0_) explained the residual species richness unexplained by clade age (Figure 4B).

## DISCUSSION

### Latitudinal comparison

Nymphalidae species richness peaks in tropical latitudes, and at least 83% of nymphalid species are found in the Neotropics, Afrotropics, Madagascar, and Southeast Asia (Figure 1). We demonstrate that higher tropical diversity does not result from simple latitudinal differences in diversification rates. Butterfly lineages in tropical regions did not consistently diversify more rapidly than in temperate regions (Figures 2 & 3). The net diversification rate in Southeast Asia was lower than in any other region except Australasia, and we found high net diversification in the Palearctic, especially during the Oligocene. Previous studies on butterflies (Papilionidae; Condamine et al. 2012) and ants (Economo et al. 2018) agree with our results and find no consistent latitudinal gradient in diversification rate, refuting the hypothesis that the LDG simply results from latitudinal differences in diversification. Instead, there is increasing evidence that the modern LDG appeared after the Eocene as a result of global climate changes throughout the Cenozoic combined with niche conservatism in tropical lineages (*e.g*., Mannion et al. 2014). We argue here that nymphalid butterflies also conform to this scenario.

The late Cretaceous climate was warmer and less seasonal (Ziegler et al. 2003). Warm and humid conditions seemed to extend to higher latitudes as documented by fossil faunas (including insects) and floras recovered in either the modern Palearctic or Nearctic (e.g. Meng & McKenna 1998). Archibald et al. (2010), for example, found Eocene insect diversity at 50° North paleolatitude to be as diverse as modern tropical diversity. According to our estimation, Nymphalidae arose around 84 Ma in present-day Eurasia and North America (Laurasia). A Laurasian origin has been reported in many different groups, such as the butterfly family Papilionidae (Condamine et al. 2012), or palm trees (Baker & Couvreur 2013). Lineage-area frequency through time shows that until the Eocene, diversity was more evenly distributed between high and low latitudes. These results suggest that nymphalid butterflies (and perhaps all butterflies) were ancestrally adapted to tropical climates and readily dispersed across tropical regions during their earliest period of diversification.

Earth’s climate cooled abruptly during the Eocene-Oligocene transition (Liu et al. 2009). The appearance of the Antarctic Circumpolar Current strengthened climatic gradients leading to more pronounced seasonality at high latitudes (Eldrett et al. 2009). Fossil evidence indicates extirpation and contraction of “tropical-like” faunas and floras towards equatorial latitudes (Saupe et al. 2019, Meseguer & Condamine 2019) and concomitant ecological turnover (Meng & McKenna 1998). We found that the contributions of the Nearctic and Palearctic fauna to global nymphalid lineage diversity were lowest during the Eocene (Figure 2). We also found that, despite being colonized shortly after the family evolved, there was a high extinction rate in the Nearctic from the early Oligocene until the mid-Miocene, probably explaining why the Nearctic region only accounts for about 3.5% of extant nymphalid diversity. Hence, early nymphalids probably occupied high latitudes of the Nearctic and eastern Palearctic until the end of the Eocene, but local extirpations and southward contractions accompanying global climate changes prevented lineages from persisting and diversifying in these regions.

Nevertheless, we found an increase in net diversification rate in the Palearctic at the end of the Eocene, and the estimated average diversification rate in the Palearctic was higher than all other regions during the Oligocene. This peak of diversification may have resulted from the emergence of cold-tolerant lineages triggered by the cooler Oligocene climate, as suggested, for example by Hawkins & DeVries (2009). This peak of diversification may also have resulted from colonization of the western Palearctic after the Turgai Sea retreated by the end of the Eocene, as suggested for other taxonomic groups (*e.g*., Mayr 2011). Our study also indicates a decrease in net diversification rate during the mid-Miocene climatic optimum before it increased again during the recent Earth cooling (Veizer & Prokoph 2015). This suggests that temperate-adapted lineages diversified faster when climate was cool and slower during warming events. We found that the disparities in Palearctic species richness were explained by recent diversification rate after removing the effect of time, thus suggesting the importance of recent events that perhaps included glaciations.

### Longitudinal comparisons

Our results identify an important role of time-for-speciation, global climate change, and the phylogenetic conservatism of ancestral tropical climatic niches as explanations for modern nymphalid LDG. However, this generalization hides great disparities in the evolutionary histories of tropical regions, each characterized by unique speciation, extinction, and dispersal histories through time (Figure 3).

Southeast Asia was central for diversification until the end of the Eocene and can be seen an ancient “cradle of diversity” (see Supplementary Information VI for an animation of the historical biogeography of nymphalid butterflies). The region seems to have been home to most of the Paleocene diversification and was a major source for lineages that dispersed to the Neotropics, Afrotropics, Palearctic, and Australasia. However, net diversification in the region greatly decreased over time relative to the other regions.

In parallel, speciation and extinction rates increased during the Miocene in the Afrotropics (including Madagascar), resulting in the highest net diversification rate among all regions during that period. African clades are generally younger than Neotropical or Asian ones, and most African clades originated during the Miocene diversification peak. The Miocene is characterized by more dispersal events from Africa towards Madagascar, which triggered speciation and multiple endemic radiations (*e.g*., *Heteropsis*, Aduse-Poku et al. 2015). More importantly, Africa experienced major climate and paleoenvironmental changes throughout the Miocene (Zhang et al. 2014). The end of the mid-Miocene climatic optimum initiated a shift from a warm and humid climate associated with trans-African forests to dry (arid) conditions, accompanied by the expansion of savannahs and C_4_ plants (Feakins et al. 2013, Jacobs 2004), probably leading to substantial species turnover.

The Neotropics are currently the most species-rich region and home to at least 37% of extant nymphalid species. Unlike Africa and Southeast Asia, which frequently exchanged lineages, we found that the Neotropics became increasingly isolated over time. We did not detect a particularly high average speciation rate in the Neotropics, except during the Eocene. Using a deep-time palynological series of Neotropical plants, Jaramillo et al. (2006) found that the rate of speciation (and total diversity) peaked during the Eocene, and probably resulted from global warming and the expansion of tropical lineages into higher latitudes. We found that the average extinction rate was remarkably low throughout the history of Neotropical diversification and that Neotropical clades tended to be older than African, Palearctic, and Nearctic clades. Therefore, our results suggest that the combination of early colonizations of the Neotropics and low extinction rates lead to the steady accumulation of lineages over time, thus supporting the hypothesis that the Neotropics are a “museum of diversity” (Stebbins 1974).

Finally, our results indicate that the extant Australasian fauna largely results from multiple dispersal events from Southeast Asia rather than local diversification. Indeed, we found a clear pattern of decreasing net diversification rate through time, but a strong increase in dispersal events during the Miocene from Southeast Asia (*ca*. 25% of global dispersal events during this period).

### Conclusion

By generating a new, large and densely sampled phylogeny of the most diverse butterfly family, the brush-footed butterflies, we showed that the history of the Nymphalidae shares fundamental similarities with other groups, including a strong LDG, a Laurasian origin, a conserved ancestral tropical niche, high Neotropical diversification during the Eocene, and higher diversification in the Palearctic during the Oligocene. However, we also unveiled notable difference with previous LDG studies, dynamics of dispersal and diversification in particular greatly varied through time and across tropical regions, in contrast to the idea that (on the evidence of this group at least) the LDG resulted from homogeneous diversification processes across all tropical regions.

## MATERIAL AND METHODS

### Time-calibrated tree

We inferred a time-calibrated tree of Nymphalidae butterflies using sequence data from the 2,866 species for which there was at least one of these 11 focal gene regions: COI, ArgKin, CAD, DDC, EF1a, GAPDH, IDH, MDH, RpS5, RpS2, wingless (Supplementary Information I). This represents ∼45% of the estimated number of species in the family. These sequence data were compiled from published and unpublished studies and subjected to multiple cleaning and verification steps. Most of the data were generated with PCR and Sanger sequencing using primers and laboratory conditions described in Wahlberg and Wheat (2008).

We generated the final tree using a tree grafting procedure (Supplementary Information I & II). First, we built a backbone tree, based on a dataset of 789 species of Nymphalidae with least six gene regions available and 11 outgroups. The topology for this backbone was generated with RAxML 8.2.12 (Stamatakis 2008) and time-calibrated using BEAST 1.8.3 (Drummond et al. 2012) using a set of 20 secondary calibrations from a recent genus-level, time-calibrated tree of all butterflies (Chazot et al. 2019). Then, we built species-level trees for 15 subclades, which often corresponded with nymphalid. For these trees, we included two outgroups and as many species as possible regardless of the amount of molecular information available. We used PartitionFinder 2.1.1 (Lanfear et al. 2017) to select partitioning strategies and substitution models for each subclade. We used BEAST 1.8.3 (Drummond et al. 2012) to estimate the topology and the relative divergence times for these subclades. Finally, the subclade trees were rescaled using the age of the root estimated by the backbone analysis, and grafted onto the backbone tree. This process was performed on the posterior distributions of both the backbone and the subclades to build a posterior distribution of 1000 grafted trees. We used TreeAnnotator 1.8.3 (Drummond et al. 2012) to summarize the tree topology with median node age and 95% credibility interval of each node. Outgroups were removed and the resulting tree was used for all subsequent analyses.

### Estimation of speciation and extinction rate

We estimated the temporal dynamics of speciation and extinction rates across our phylogeny using BAMM 2.5 (Rabosky et al. 2013; Rabosky 2014; Rabosky et al. 2017; Supplementary Information III). We accounted for missing species by specifying the sampling fraction at the genus level. The analyses were run for 50 million generations with four reversible-jump MCMC, sampling parameters every 50,000 generations. The output was then analysed using the R package BAMMtools. We checked that the MCMC converged with effective sample size above 600 after we discarded the first 10% of samples as burn-in.

### Inference of biogeographic history

We performed a maximum-likelihood estimate of geographic range evolution using the Dispersal-Extinction-Cladogenesis (DEC) model (Ree & Smith 2008) as implemented in an extended C++ version of DEC (Smith 2009), called DECX (Beeravolu & Condamine 2016). We designed a biogeographic model spanning the evolutionary history of Nymphalidae, starting in the Late Cretaceous (Supplementary Information IV). We assigned extant species to nine biogeographic regions: western Nearctic, eastern Nearctic, western Palearctic, eastern Palearctic, Neotropics, Afrotropics, India, Southeast Asia, and Australasia. We designed a time-stratified model in which both the adjacency matrices and dispersal matrices varied between time periods. Time was divided into five time periods: 100-80, 80-60, 60-30, 30-10, 10-0 Myr to account for increasing or decreasing connectivity between biogeographic regions through time.

### Biogeographic patterns of diversification: combining BAMM and DEC

We combined the speciation and extinction rate estimates for ancestral lineages obtained from BAMM with the biogeographic ranges and timing of dispersal events estimated with DECX to compare the accumulation of dispersal events, the average speciation rates, and the average extinction rates among regions (Supplementary Information V).

#### Dispersal events

We identified the range with the highest probability at each node. For each dispersal event, we drew 1,000 random times of dispersal events along the branches. For each replicate, we recorded the number of dispersal events between biogeographic regions occurring during four geological time periods transformed these sums of events into percentages of the total number of events during that time period. For this analysis we reduced the number of areas from nine to six by combining eastern and western Nearctic into Nearctic, eastern and western Palearctic into Palearctic, India and Southeast Asia into Southeast Asia; Australasia was kept as a single area (Supplementary Information V).

#### Lineage-area frequency through time

We used DECX results to estimate the frequency of lineages sampled in the tree in each area through time. The number of lineages was computed within 0.5 Myr time intervals. If a dispersal event occurred along a branch, we assumed that it occurred at the branch midpoint.

#### Biogeography and diversification rates

We estimated variation in speciation and extinction rates through time within each biogeographic region by combining DECX ancestral range estimates and speciation/extinction rates estimation from BAMM (Supplementary Information V). We recovered the rates speciation and extinction rates through time for all branches using the function dtRates (BAMMtools, Rabosky et al. 2014). We used a sliding window analysis to estimate the mean diversification rates through time for each biogeographic region. We computed the average speciation, extinction, and net diversification rates per region within 4 Myr time windows by shifting the window by 1 Myr. Within each time window, if a lineage occupied an area, the rates estimated for this branch (or fraction of the branch in case of dispersal events) contributed to the average rate of the region. The average was computed by estimating the number of events in one area (rate*branch length) divided by the sum of branch lengths occupying this area during the same time-interval. We repeated the analysis for 100 random timings of dispersal events.

#### Animated historical biogeography

To help visualizing the pattern of historical biogeography we displayed a single realization of historical dispersal events through time on a map with present-day positions of continents for simplicity of implementation. We also displayed at the same time the average net diversification rate through time in regions and the relative frequency of lineages in different regions through time (Supplementary Information VI). The code was adapted from Dudas et al. (2017).

### Biogeographic patterns of diversification: regional diversification

We investigated diversification by analysing clades that diversified within a single region (Supplementary Information VII). We refer to such events “regional diversification”. We arbitrarily defined a regional diversification event as a clade of at least four terminals (*i.e*., extant taxa included in our tree) that has diversified in a single biogeographic region. We circumscribed fewer regions for this approach: Neotropics, Afrotropics, Southeast Asia combined with Australasia, Palearctic, Nearctic because of the large number of dispersal events happening between some of the areas. We identified 90 local diversification events, which represent an estimated 5373 total species (*ca*. 86% of all nymphalid diversity). Using these 90 cases, we investigated whether (1) any pattern of diversification emerges among regional diversification events within the same biogeographic region, for example in terms of time variation of speciation rate and whether (2) the age of the crown, recent dynamics of diversification or dynamics occurring during the early stage of diversification explains the extant diversity of these radiations.

For each regional diversification event, we fitted a model in which speciation rate was modelled as an exponential function of time, while extinction rate remained constant (Morlon et al. 2011) using the R-package RPANDA 1.5 (Morlon et al. 2016). For each clade, we estimated the extinction parameter (μ) and the two parameters for speciation: speciation rate at present (λ_0_) and coefficient of time variation (α). Using these parameters, we also computed the speciation rate at the crown age (λ_crown_), the net diversification at present (netDiv_0_), and the net diversification rate at the crown age (netDiv_crown_). We tested for a relationship between crown age, netDiv_0_, netDiv_crown_, or α and the estimated total number of extant species (log-transformed species richness) by fitting a linear model for each biogeographic region (Supplementary Information VII). Since crown age had a strong and significant effect on the extant number of species, we performed again the linear models for each region on the residual species richness after removing the effect of crown age.

## Supporting information

Figure SIII. 1

Figure SIV. 1

## Acknowledgements

DJL was supported by grants DEB-1541557 from NSF and WW-227R-17 from the National Geographic Society. RIVW was supported by Leverhulme Trust emeritus programme. ME was supported by an ATIP grant, a grant from the Human Frontier Science Program (RGP0014/2016) and a grant from the French National Research Agency (ANR CLEARWING ANR-16-CE02-0012). EOA was supported by Sigma-Xi (G20100315153261), Center for Systematic Entomology and the Council of the Linnean Society and the Systematics Association for the Systematics Research Fund. SN was supported by the Swedish Research Council (2015-04218 and 2019-03441).

