## Supplementary Information for "The latitudinal diversity gradient in brush-footed butterflies (Nymphalidae): conserved ancestral tropical niche but different continental histories"

### I - Gene and taxon sampling

We based our analyses on 11 gene regions: the mitochondrial COI gene (1473 bp) and the nuclear genes ArgKin (596 bp), CAD (850 bp), DDC (373 bp), EF1a (1240 bp), GAPDH (691 bp), IDH (710 bp), MDH (733 bp), RpS5 (617 bp), RpS2 (411 bp), and wingless (412 bp). These gene regions have been used extensively in molecular systematic studies of Nymphalidae. Most unpublished sequences that we use have been generated according to protocols reported in Wahlberg and Wheat (2008).

We assembled our dataset from two databases: our own VoSeq (Peña and Malm 2012) and BoLD (Ratnasingham & Hebert 2007). From each of these databases we extracted all the individuals classified as "Nymphalidae", which gave us 10,477 individuals and 23,138 individuals respectively (as of May 2016). In the VoSeq dataset, we filtered multiple hits of species names by keeping only one species representative having the highest number of base pairs available. This first list of taxa was compared to the dataset imported from BoLD, which allowed adding 415 taxa absent from VoSeq for a total of 3,070 taxa. This list of taxa was further cleaned by checking for misspellings of names leading to multiple representatives per species. We also removed all the barcode sequences imported from BoLD that were not associated with a published paper on GenBank. We checked for the validity of all species names, mostly relying on online resources (e.g. http://ftp.funet.fi/pub/sci/bio/life/insecta/lepidoptera/ditrysia/papilionoidea/nymphalidae/). When we encountered a disagreement on taxon assignment we searched for the most recent relevant source of information to decide whether the taxon was included as a valid species or removed. All these sequences were aligned in Codon Code v.7.1.2 (CodonCode Corporation, www.codoncode.com) and uploaded into our VoSeq database. Finally, we used RAxML v.8.2.12 (Stamatakis 2008) on the full dataset, to search for the best scoring topology. We partitioned the dataset into 4 subsets: 1$^{st}$ position of all nuclear genes, 2$^{nd}$ position of all nuclear genes, 3$^{rd}$ position of all nuclear genes, 1$^{st}$ and 2$^{nd}$ position of COI (3$^{rd}$ position of COI was excluded). We selected a GTR+G to model substitution rates for each partition. This RAxML analysis was used to identify misidentifications, rogue taxa, and other species with unexpected phylogenetic positions likely resulting from human/laboratory error and to check for misalignments. Rogue taxa were removed from the dataset. After these cleaning steps, we ended up with a final taxon sampling of 2,866 species (Table SI. 1).

### II - Time-calibrated phylogeny

#### *1 - Overview*

The scale of our dataset prevented us from a direct timing of divergence analysis of the full dataset. Therefore, we adopted a tree grafting procedure (e.g., Jetz et al. 2012, Tonini et al. 2016, Rabosky et al. 2018). We first built a backbone tree, based on a dataset of 800 taxa with at least 6 sequenced gene regions available. This backbone was time-calibrated using a set of 20 secondary calibrations from a recent genus level time-calibrated tree of all Papilionoidea (Chazot et al. 2019). We then inferred 15 species-level trees, corresponding often to the different subfamilies. For these trees, we included as many species as possible regardless of the amount of molecular information available, and we estimated relative times of divergence. Finally, these subclade trees were rescaled using the age of the root estimated in the backbone analysis, to be grafted on the backbone tree. This process was performed on the posterior distributions of both the backbone and the subclades to build a posterior distribution of grafted trees, each tree with 2,866 species.

#### 2 - *Backbone dataset*

To build a robust time-calibrated backbone tree, we created a subset of taxa that had at least 6 gene regions out of 11 to limit the potential effect of missing information in our molecular dataset. All genera were represented in this subset, which contained 1,172 taxa. We further refined this list by removing some taxa from clades that were overrepresented. As such, our final dataset contained 789 species classified as Nymphalidae. We added 11 outgroups, including Lycaenidae and Riodinidae – the two sister families of Nymphalidae - and Pieridae, sister family to Lycaenidae + Riodinidae + Nymphalidae. Preliminary analyses revealed potential saturation with the 3$^{rd}$ codon position of COI for the time-calibration analysis. Thus, we excluded this position from the backbone analyses.

#### *3 - Time-calibrated backbone phylogeny*

For the backbone tree, we followed Chazot et al. (2019)'s partitioning strategy for the backbone: 1$^{st}$ position of all nuclear genes, 2$^{nd}$ position of all nuclear genes, 3$^{rd}$ position of all nuclear genes, 1$^{st}$ and 2$^{nd}$ position of COI. For each subset, the substitution rate was modelled by a GTR+G. First, we performed a maximum likelihood search with RAxML v.8.2.12 (Stamatakis 2008) to obtain a topology. This topology was almost identical to that obtained in Chazot et al. (2019)'s time calibration of butterflies, which is our source of secondary calibrations for this study. To ensure correct estimation of the crown and stem ages of the subclades that were to be grafted, we compared the subclade tree topologies (see below), and checked that the deepest node within each subclade was included. This backbone topology was then time-calibrated using BEAST 1.8.3. (Drummond et al 2012). To time-calibrate the backbone tree we used the same partitioning strategy as for RAxML and GTR+G to model the rates of substitution. For each subset, we set an uncorrelated lognormal relaxed clock (unlinked across partitions, four independent clocks). For calibrating the clocks, we relied on secondary calibrations taken from Chazot et al. (2019). The butterfly fossil record is depauperate, which drastically limits their use in time-calibrations (de Jong 2017). Chazot et al. (2019) estimated a time-calibrated genus-level tree of butterflies, using the most complete set of fossils up to date, which provides the most robust age estimates for the root of Nymphalidae and internal relationships to date. Using these estimations, we defined a set of 20 calibration points across the tree (Table SII. 1). We used normal priors for these calibration points, where the mean and standard deviation were adjusted to recover the age and credibility interval from Chazot et al. (2019). We ran eight independent BEAST analyses for 40 million generations each, sampling every 40,000 generations. We checked for convergence of the Markov chains Monte Carlo (MCMC) using Tracer v.1.6. After removing the first 10 million generations as burn-in we combined the different runs using LogCombiner 1.8.3 (Drummond et al 2012). Posterior distributions of node ages were synthesized by extracting the median and 95% credibility interval (CI) for each node. This tree was compared to that of Chazot et al (2019) to ensure that we recovered expected ages.

**Table SII. 1.** Clades corresponding to the nodes constrained for time-calibrating the backbone tree. Information about the shape of normal prior (mean and standard deviation) is also given.

| Clade | mean | stdev |
|---|---|---|
| root | 83 | 120 |
| Apaturinae | 33.6 | 4.00 |
| Biblidinae | 40.1 | 4.00 |
| Charaxinae | 41 | 4.0 |
| Cyrestinae | 33.4 | 4.0 |
| Danainae | 42 | 4 |
| Heliconiinae | 47.7 | 4.5 |
| Limenitidinae | 48.7 | 4.5 |
| Nymphalidae | 82 | 7.5 |
| Nymphalinae | 47.5 | 4.5 |
| Satyrinae | 54.4 | 4.5 |

| | | |
|---|---|---|
| Ithomiini | 25.6 | 2.8 |
| Argynnini | 29.2 | 3.0 |
| Adoliadini | 32.2 | 3.2 |
| Limenitidini | 16.3 | 2 |
| Nymphalini | 35 | 3.5 |
| Melitaeini | 27.4 | 2.5 |
| Charaxini_Euxanthini | 17.5 | 2.2 |
| Mycalesina | 24.2 | 2.2 |
| Satyrini | 37.2 | 3.5 |

*4 - Subclade trees*

We subdivided the whole family Nymphalidae into 15 monophyletic groups of species (sublades), all well defined by previous literature. These subclades corresponded to the Danaini, Ithomiini, Heliconiinae, Cyrestinae, Charaxinae, Libythaeinae, Nymphalinae, Apaturinae, Biblidinae, Calinaginae, Limenitidinae, Pseudergolinae, Satyrinae group I (*Heteropsis*+*Coenonympha*), Satyrinae group II (*Elymnias*+*Morpho*) and Satyrini. For each of them we included all the species available. We also added two outgroups, one from the sister clade of the focal subclade and one sister clade to the two previous ones. For each subclade dataset we started by assessing the best partitioning strategy using PartitionFinder 2 (Lanfear et al. 2017), allowing all possible combinations of genes and codon position and all substitution models available in BEAST. Partitioning strategies were identified using the *greedy* algorithm and BIC score. By default, we started by setting one molecular clock (uncorrelated relaxed clock with lognormal distribution) for each partition. If convergence or good mixing could not be obtained after running BEAST, we reduced the number of molecular clocks (see detailed description of subclade analyses). We did not add any time-calibration and therefore only estimated the relative divergence times. We enforced the monophyly of the clade to be grafted (i.e., excluding the outgroups); all other relationships were estimated by BEAST. We performed at least two independent runs with BEAST for each subclade. The length of the MCMC varied according to the size of the subclade (see detailed description of subclade analyses descriptions). We checked for convergence and mixing of the MCMC using Tracer v.1.6.0 (Rambaut et al 2014). Posterior distributions were finally combined after removing an appropriate burn-in fraction.

Details of the analysis for each subclade are provided hereafter.

a) Apaturinae

*Dataset* – The dataset for the Apaturinae consisted of 32 taxa to which two outgroups were added: *Marpesia eleuchea* (Cyrestinae), *Biblis hyperia* (Biblidinae).

*Partition Finder* – Partition Finder identified 10 subsets.

| Subset # | Substitution model | Gene fragments and codon positions |
|---|---|---|
| Subset 1 | GTR+I+G | MDH_pos1, IDH_pos1, CAD_pos1, GAPDH_pos1, ArgKin_pos1, RpS2_pos1, RpS5_pos1, EF1a_pos1 |
| Subset 2 | TRN+I | GAPDH_pos2, RpS5_pos2, EF1a_pos2, ArgKin_pos2, MDH_pos2, CAD_pos2, IDH_pos2 |
| Subset 3 | GTR+G | wingless_pos3, ArgKin_pos3, GAPDH_pos3, EF1a_pos3 |
| Subset 4 | HKY+I | CAD_pos3 |
| Subset 5 | HKY+G | COI-begin_pos3, COI-end_pos3 |
| Subset 6 | GTR+G | COI-begin_pos1, COI-end_pos1 |
| Subset 7 | HKY+I | COI-begin_pos2, COI-end_pos2 |
| Subset 8 | GTR+G | DDC_pos3, RpS2_pos3, RpS5_pos3, IDH_pos3, MDH_pos3 |
| Subset 9 | K80 | DDC_pos1, wingless_pos2, wingless_pos1 |
| Subset 10 | JC | RpS2_pos2, DDC_pos2 |

*BEAST analysis* – In order to improve the quality of our runs we replaced the default priors for rates of substitutions by uniform prior ranging between 0 and 10 for the following cases: subset1.at, cg, gt, subset6.ac, at, cg, gt. We used a single molecular clock linked across all partition to obtain good mixing and convergence. We used a birth-death tree prior. We performed two runs of 60 million generations, sampling trees and parameters every 6000 generations.

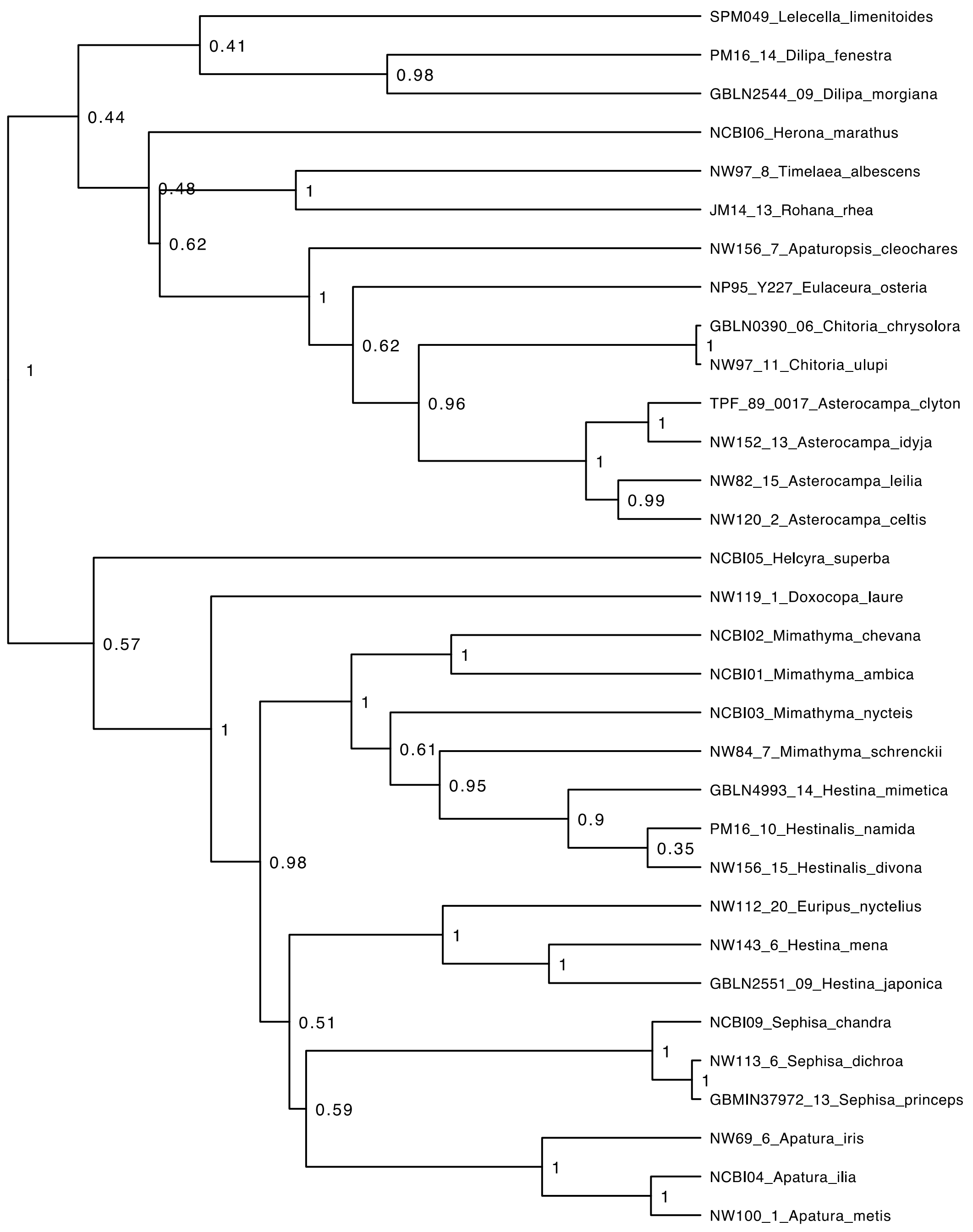


**Figure SII. 1.** Maximum clade credibility tree for the Apaturinae subclade from BEAST. Posterior probabilities are indicated at the nodes.

b) Biblidinae

*Dataset* – The dataset for the Biblidinae consisted of 155 taxa to which two outgroups were added: *Marpesia eleuchea* (Cyrestinae), *Chitoria ulupi* (Apaturinae).

*Partition Finder* – Partition Finder identified 12 subsets.

| Subset # | Substitution model | Gene fragments and codon positions |
|---|---|---|
| Subset 1 | GTR+I+G | DDC_pos1, ArgKin_pos1, RpS2_pos1, MDH_pos1, RpS5_pos1, IDH_pos1, wingless_pos1, CAD_pos1 |
| Subset 2 | GTR+I+G | COI-begin_pos2, EF1a_pos2, GAPDH_pos2, RpS5_pos2, ArgKin_pos2, DDC_pos2, IDH_pos2, CAD_pos2, MDH_pos2, COI-end_pos2 |
| Subset 3 | GTR+G | ArgKin_pos3 |
| Subset 4 | HKY+I+G | MDH_pos3, CAD_pos3 |
| Subset 5 | GTR+I+G | COI-begin_pos3 |
| Subset 6 | GTR+I+G | COI-end_pos1, COI-begin_pos1 |
| Subset 7 | HKY+G | COI-end_pos3 |
| Subset 8 | GTR+I+G | DDC_pos3, IDH_pos3, GAPDH_pos3 |
| Subset 9 | GTR+G | EF1a_pos3 |
| Subset 10 | TRN+I+G | GAPDH_pos1, EF1a_pos1 |
| Subset 11 | JC+G | RpS2_pos2, wingless_pos2 |
| Subset 12 | HKY+G | wingless_pos3, RpS5_pos3, RpS2_pos3 |

*BEAST analysis* – In order to improve the quality of our runs we replaced the default priors for rates of substitutions by uniform prior ranging between 0 and 10 for the following cases: subset3.cg, gt, subset5.at, ag, cg. We used one molecular clock per subset identified by Partition Finder and obtained good mixing and convergence. We used a birth-death tree prior. We performed two runs of 150 million generations, sampling trees and parameters every 15000 generations.

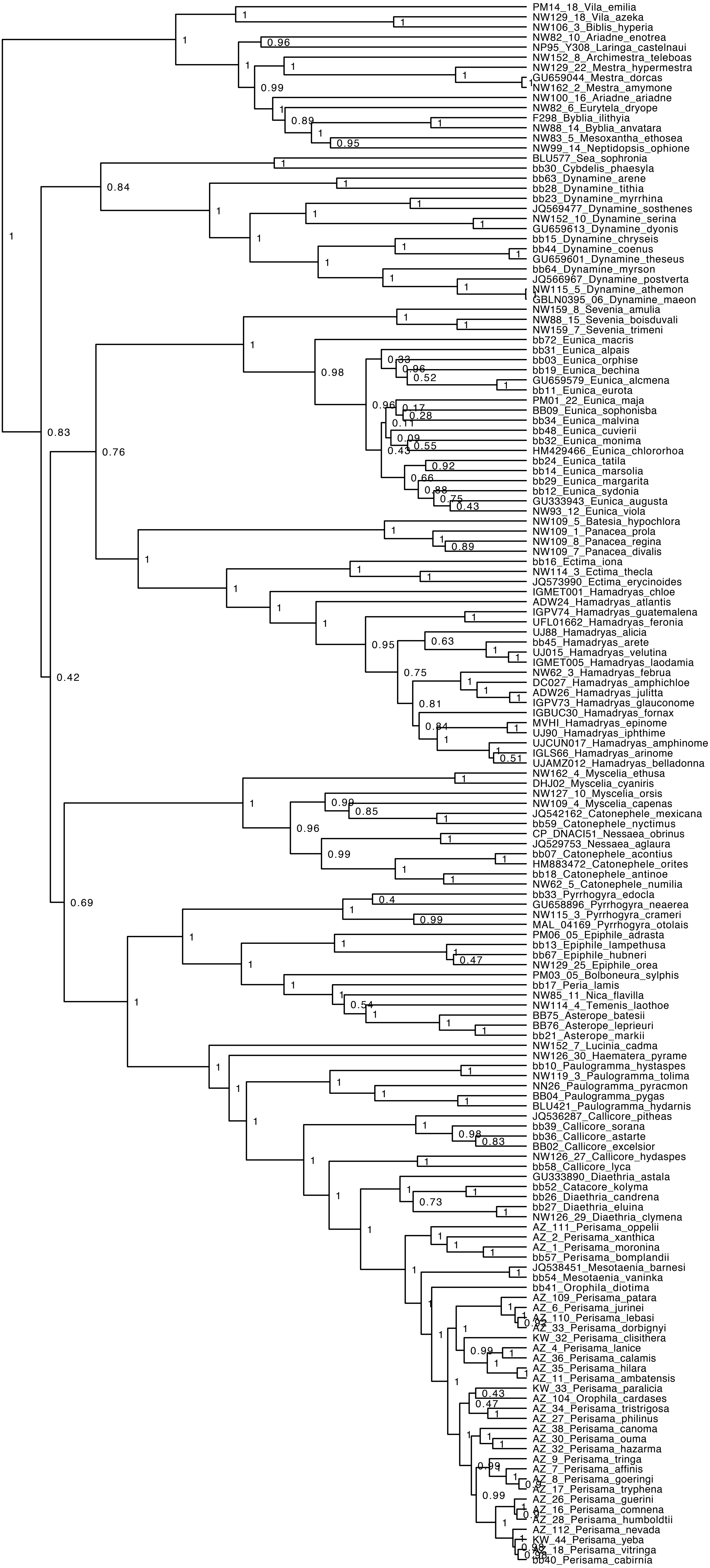

PM14_18_Vila_emilia
NW129_18_Vila_azeka
NW106_3_Biblis_hyperia
NW82_10_Ariadne_enotrea
NP95_Y308_Laringa_castelnaui
NW152_8_Archimestra_teleboas
NW129_22_Mestra_hypermestra
GU659044_Mestra_dorcas
NW162_2_Mestra_amymone
NW100_16_Ariadne_ariadne
NW82_6_Eurytela_dryope
F298_Byblia_ilithyia
NW88_14_Byblia_anvatara
NW83_5_Mesoxantha_ethosea
NW99_14_Neptidopsis_ophione
BLU577_Sea_sophronia
bb30_Cybdelis_phaesyla
bb63_Dynamine_arene
bb28_Dynamine_tithia
bb23_Dynamine_myrrhina
JQ569477_Dynamine_sosthenes
NW152_10_Dynamine_serina
GU659613_Dynamine_dyonis
bb15_Dynamine_chryseis
bb44_Dynamine_coenus
GU659601_Dynamine_theseus
bb64_Dynamine_myrson
JQ566967_Dynamine_postverta
NW115_5_Dynamine_athemon
GBLN0395_06_Dynamine_maeon
NW159_8_Sevenia_amulia
NW88_15_Sevenia_boisduvali
NW159_7_Sevenia_trimeni
bb72_Eunica_macris
bb31_Eunica_alpais
bb03_Eunica_orphise
bb19_Eunica_bechina
GU659579_Eunica_alcmena
bb11_Eunica_eurota
PM01_22_Eunica_maja
BB09_Eunica_sophonisba
bb34_Eunica_malvina
bb48_Eunica_cuvierii
bb32_Eunica_monima
HM429466_Eunica_chlororhoa
bb24_Eunica_tatila
bb14_Eunica_marsolia
bb29_Eunica_margarita
bb12_Eunica_sydonia
GU333943_Eunica_augusta
NW93_12_Eunica_viola
NW109_5_Batesia_hypochlora
NW109_1_Panacea_prola
NW109_8_Panacea_regina
NW109_7_Panacea_divalis
bb16_Ectima_iona
NW114_3_Ectima_thecla
JQ573990_Ectima_erycinoides
IGMET001_Hamadryas_chloe
ADW24_Hamadryas_atlantis
IGPV74_Hamadryas_guatemalena
UFL01662_Hamadryas_feronia
UJ88_Hamadryas_alicia
bb45_Hamadryas_arete
UJ015_Hamadryas_velutina
IGMET005_Hamadryas_laodamia
NW62_3_Hamadryas_februa
DC027_Hamadryas_amphichloe
ADW26_Hamadryas_julitta
IGPV73_Hamadryas_glauconome
IGBUC30_Hamadryas_fornax
MVHI_Hamadryas_epinome
UJ90_Hamadryas_iphthime
UJCUN017_Hamadryas_amphinome
IGLS66_Hamadryas_arinome
UJAMZ012_Hamadryas_belladonna
NW162_4_Myscelia_ethusa
DHJ02_Myscelia_cyaniris
NW127_10_Myscelia_orsis
NW109_4_Myscelia_capenas
JQ542162_Catonephele_mexicana
bb59_Catonephele_nyctimus
CP_DNACI51_Nessaea_obrinus
JQ529753_Nessaea_aglaura
bb07_Catonephele_acontius
HM883472_Catonephele_orites
bb18_Catonephele_antinoe
NW62_5_Catonephele_numilia
bb33_Pyrrhogyra_edocla
GU658896_Pyrrhogyra_neaerea
NW115_3_Pyrrhogyra_crameri
MAL_04169_Pyrrhogyra_otolais
PM06_05_Epiphile_adrasta
bb13_Epiphile_lampethusa
bb67_Epiphile_hubneri
NW129_25_Epiphile_orea
PM03_05_Bolboneura_sylphis
bb17_Peria_lamis
NW85_11_Nica_flavilla
NW114_4_Temenis_laothoe
BB75_Asterope_batesii
BB76_Asterope_leprieuri
bb21_Asterope_markii
NW152_7_Lucinia_cadma
NW126_30_Haematera_pyrame
bb10_Paulogramma_hystaspes
NW119_3_Paulogramma_tolima
NN26_Paulogramma_pyracmon
BB04_Paulogramma_pygas
BLU421_Paulogramma_hydarnis
JQ536287_Callicore_pitheas
bb39_Callicore_sorana
bb36_Callicore_astarte
BB02_Callicore_excelsior
NW126_27_Callicore_hydaspes
bb58_Callicore_lyca
GU333890_Diaethria_astala
bb52_Catacore_kolyma
bb26_Diaethria_candrena
bb27_Diaethria_eluina
NW126_29_Diaethria_clymena
AZ_111_Perisama_oppelii
AZ_2_Perisama_xanthica
AZ_1_Perisama_moronina
bb57_Perisama_bomplandii
JQ538451_Mesotaenia_barnesi
bb54_Mesotaenia_vaninka
bb41_Orophila_diotima
AZ_109_Perisama_patara
AZ_6_Perisama_jurinei
AZ_110_Perisama_lebasi
AZ_33_Perisama_dorbignyi
KW_32_Perisama_clisithera
AZ_4_Perisama_lanice
AZ_36_Perisama_calamis
AZ_35_Perisama_hilara
AZ_11_Perisama_ambatensis
KW_33_Perisama_paralicia
AZ_104_Orophila_cardases
AZ_34_Perisama_tristrigosa
AZ_27_Perisama_philinus
AZ_38_Perisama_canoma
AZ_30_Perisama_ouma
AZ_32_Perisama_hazarma
AZ_9_Perisama_tringa
AZ_7_Perisama_affinis
AZ_8_Perisama_goeringi
AZ_17_Perisama_tryphena
AZ_26_Perisama_guerini
AZ_16_Perisama_comnena
AZ_28_Perisama_humboldtii
AZ_112_Perisama_nevada
KW_44_Perisama_yeba
AZ_18_Perisama_vitringa
bb40_Perisama_cabirnia

**Figure SII. 2.** (page before). Maximum clade credibility tree for the Biblidinae subclade from BEAST. Posterior probabilities are indicated at the nodes.

c) Calinaginae

*Dataset* – The dataset for the Calinaginae consisted of 6 taxa to which two outgroups were added: *Heliconius melpomene* (Heliconiinae), *Amathusia phidippus* (Satyrinae).

*Partition Finder* – Partition Finder identified 9 subsets.

| Subset # | Substitution model | Gene fragments and codon positions |
|---|---|---|
| Subset 1 | GTR | ArgKin_pos1, RpS2_pos1, EF1a_pos1, wingless_pos1, DDC_pos1, RpS5_pos1, GAPDH_pos1, CAD_pos1, MDH_pos1, IDH_pos1 |
| Subset 2 | HKY | RpS5_pos2, MDH_pos2, GAPDH_pos2, DDC_pos2, ArgKin_pos2, IDH_pos2, EF1a_pos2 |
| Subset 3 | GTR | EF1a_pos3, ArgKin_pos3 |
| Subset 4 | TRN | MDH_pos3, CAD_pos3 |
| Subset 5 | TRN+I | CAD_pos2, COI-end_pos1, COI-begin_pos1 |
| Subset 6 | HKY+I | COI-end_pos3, COI-begin_pos3 |
| Subset 7 | HKY | COI-end_pos2, COI-begin_pos2 |
| Subset 8 | GTR | RpS2_pos3, IDH_pos3, RpS5_pos3, GAPDH_pos3, DDC_pos3, wingless_pos3 |
| Subset 9 | JC | wingless_pos2, RpS2_pos2 |

*BEAST analysis* – In order to improve the quality of our runs, we replaced the default priors for rates of substitutions by uniform prior ranging between 0 and 10 for the following cases: subset3.cg, subset8.cg, subset1.at, gt. We used a single molecular clock linked across all partition to obtain good mixing and convergence. We used a birth-death tree prior. We performed two runs of 50 million generations, sampling trees and parameters every 5000 generations.

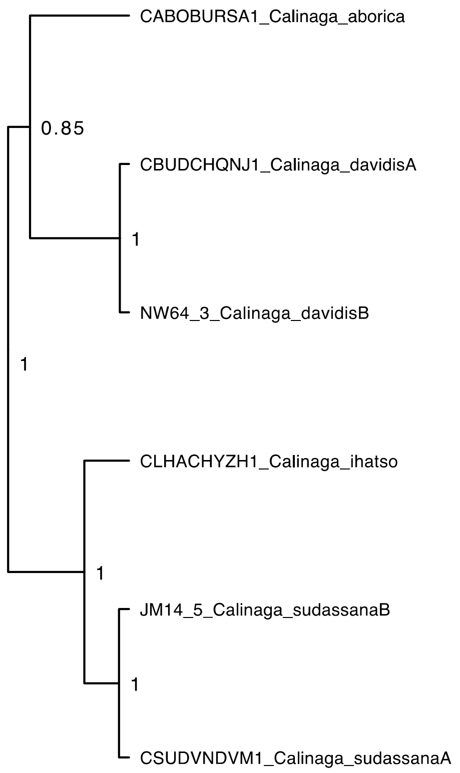


**Figure SII. 3.** Maximum clade credibility tree for the Calinaginae subclade from BEAST. Posterior probabilities are indicated at the nodes.

d) Charaxinae

*Dataset* – The dataset for the Charaxinae consisted of 234 taxa to which two outgroups were added: *Calinaga davidis* (Calinaginae), *Amathusia phidippus* (Satyrinae).

*Partition Finder* – Partition Finder identified 11 subsets.

| Subset # | Substitution model | Gene fragments and codon positions |
|---|---|---|
| Subset 1 | TRN+I+G | ArgKin_pos1, GAPDH_pos1, MDH_pos1, EF1a_pos1, RpS2_pos1, RpS5_pos1 |
| Subset 2 | HKY+I+G | MDH_pos2, RpS5_pos2, EF1a_pos2, GAPDH_pos2, ArgKin_pos2, DDC_pos2, IDH_pos2 |
| Subset 3 | TRN | RpS2_pos2, wingless_pos2, ArgKin_pos3 |
| Subset 4 | HKY+I+G | MDH_pos3, CAD_pos3 |
| Subset 5 | GTR+G | DDC_pos1, wingless_pos1, CAD_pos1, IDH_pos1 |
| Subset 6 | TRN+I | CAD_pos2, COI-begin_pos2, COI-end_pos2 |
| Subset 7 | GTR+G | COI-end_pos3, COI-begin_pos3 |
| Subset 8 | TRN+I+G | COI-end_pos1, COI-begin_pos1 |
| Subset 9 | GTR+I+G | DDC_pos3, IDH_pos3, RpS2_pos3, GAPDH_pos3 |
| Subset 10 | GTR+I+G | EF1a_pos3, RpS5_pos3 |
| Subset 11 | K80+I+G | wingless_pos3 |

*BEAST analysis* – In order to improve the quality of our runs we replaced the substitution model of subset3 by K80. We used one molecular clock per subset identified by Partition Finder and obtained good mixing and convergence. We used a birth-death tree prior. We performed two runs of 100 million generations, sampling trees and parameters every 10000 generations.

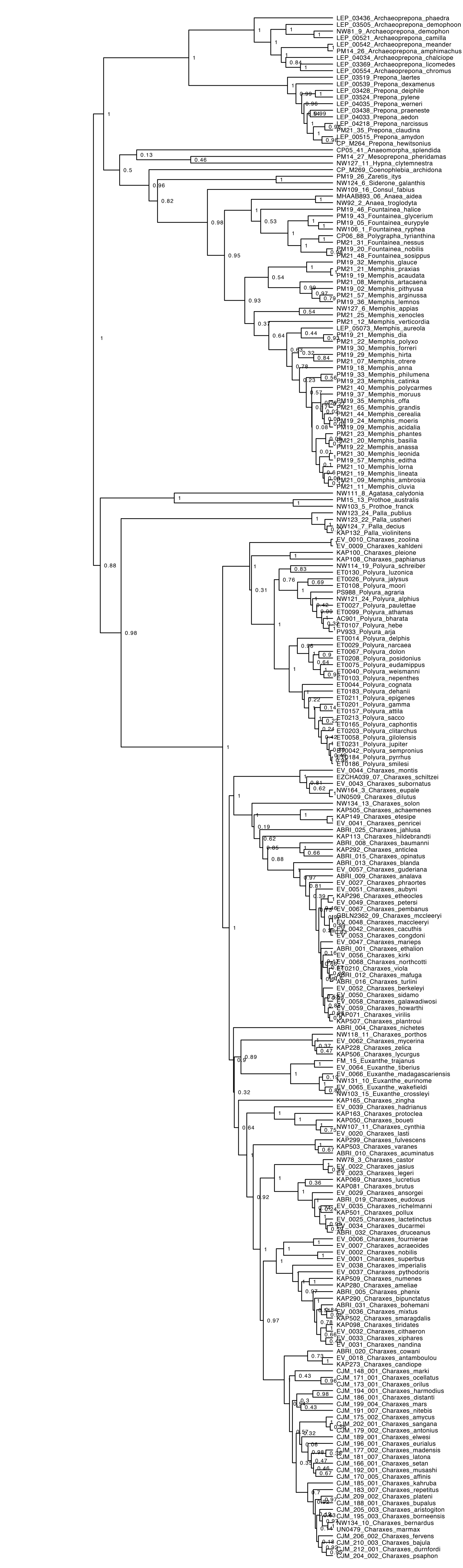
LEP_03436_Archaeoprepona_phaedra
LEP_03505_Archaeoprepona_demophoon
NW81_9_Archaeoprepona_demophon
LEP_00521_Archaeoprepona_camilla
LEP_00542_Archaeoprepona_meander
PM14_26_Archaeoprepona_amphimachus
LEP_04034_Archaeoprepona_chalciope
LEP_03369_Archaeoprepona_licomedes
LEP_00554_Archaeoprepona_chromus
LEP_03519_Prepona_laertes
LEP_00539_Prepona_dexamenus
LEP_03428_Prepona_deiphile
LEP_03524_Prepona_pylene
LEP_04035_Prepona_werneri
LEP_03438_Prepona_praeneste
LEP_04033_Prepona_aedon
LEP_04218_Prepona_narcissus
PM21_35_Prepona_claudina
LEP_00515_Prepona_amydon
CP_M264_Prepona_hewitsonius
CP05_41_Anaeomorpha_splendida
PM14_27_Mesoprepona_pheridamas
NW127_11_Hypna_clytemnestra
CP_M269_Coenophlebia_archidona
PM19_26_Zaretis_itys
NW124_6_Siderone_galanthis
NW109_16_Consul_fabius
MHAAB893_06_Anaea_aidea
NW92_2_Anaea_troglodyta
PM19_46_Fountainea_halice
PM19_43_Fountainea_glycerium
PM19_05_Fountainea_eurypyle
NW106_1_Fountainea_ryphea
CP06_88_Polygrapha_tyrianthina
PM21_31_Fountainea_nessus
PM19_20_Fountainea_nobilis
PM21_48_Fountainea_sosippus
PM19_32_Memphis_glauce
PM21_21_Memphis_praxias
PM19_19_Memphis_acaudata
PM21_08_Memphis_artacaena
PM19_02_Memphis_pithyusa
PM21_57_Memphis_arginussa
PM19_36_Memphis_lemnos
NW127_6_Memphis_appias
PM21_25_Memphis_xenocles
PM21_12_Memphis_verticordia
LEP_05073_Memphis_aureola
PM19_21_Memphis_dia
PM21_22_Memphis_polyxo
PM19_30_Memphis_forreri
PM19_29_Memphis_hirta
PM21_07_Memphis_otrere
PM19_18_Memphis_anna
PM19_33_Memphis_philumena
PM19_23_Memphis_catinka
PM21_40_Memphis_polycarmes
PM19_37_Memphis_moruus
PM19_35_Memphis_offa
PM21_65_Memphis_grandis
PM21_44_Memphis_cerealia
PM19_24_Memphis_moeris
PM19_09_Memphis_acidalia
PM21_23_Memphis_phantes
PM21_20_Memphis_basilia
PM19_22_Memphis_anassa
PM21_30_Memphis_leonida
PM19_57_Memphis_editha
PM21_10_Memphis_lorna
PM21_19_Memphis_lineata
PM21_09_Memphis_ambrosia
PM21_11_Memphis_cluvia
NW111_8_Agatasa_calydonia
PM15_13_Prothoe_australis
NW103_5_Prothoe_franck
NW123_24_Palla_publius
NW123_22_Palla_ussheri
NW124_7_Palla_decius
KAP132_Palla_violinitens
EV_0010_Charaxes_zoolina
EV_0009_Charaxes_kahldeni
KAP100_Charaxes_pleione
KAP108_Charaxes_paphianus
NW114_19_Polyura_schreiber
ET0130_Polyura_luzonica
ET0026_Polyura_jalysus
ET0108_Polyura_moori
PS988_Polyura_agraria
NW121_24_Polyura_alphius
ET0027_Polyura_paulettae
ET0099_Polyura_athamas
AC901_Polyura_bharata
ET0107_Polyura_hebe
PV933_Polyura_arja
ET0014_Polyura_delphis
ET0029_Polyura_narcaea
ET0067_Polyura_dolon
ET0208_Polyura_posidonius
ET0075_Polyura_eudamippus
ET0040_Polyura_weismanni
ET0103_Polyura_nepenthes
ET0044_Polyura_cognata
ET0183_Polyura_dehanii
ET0211_Polyura_epigenes
ET0201_Polyura_gamma
ET0157_Polyura_attila
ET0213_Polyura_sacco
ET0165_Polyura_caphontis
ET0203_Polyura_clitarchus
ET0058_Polyura_gilolensis
ET0231_Polyura_jupiter
ET0042_Polyura_sempronius
ET0184_Polyura_pyrrhus
ET0186_Polyura_smilesi
EV_0044_Charaxes_montis
EZCHA039_07_Charaxes_schiltzei
EV_0043_Charaxes_subornatus
NW164_3_Charaxes_eupale
UN0509_Charaxes_dilutus
NW134_13_Charaxes_solon
KAP505_Charaxes_achaemenes
KAP149_Charaxes_etesipe
EV_0041_Charaxes_penricei
ABRI_025_Charaxes_jahlusa
KAP113_Charaxes_hildebrandti
ABRI_008_Charaxes_baumanni
KAP292_Charaxes_anticlea
ABRI_015_Charaxes_opinatus
ABRI_013_Charaxes_blanda
EV_0057_Charaxes_guderiana
ABRI_009_Charaxes_analava
EV_0027_Charaxes_phraortes
EV_0051_Charaxes_aubyni
KAP296_Charaxes_etheocles
EV_0049_Charaxes_petersi
EV_0067_Charaxes_pembanus
GBLN2362_09_Charaxes_mccleeryi
EV_0048_Charaxes_maccleeryi
EV_0042_Charaxes_cacuthis
EV_0053_Charaxes_congdoni
EV_0047_Charaxes_marieps
ABRI_001_Charaxes_ethalion
EV_0056_Charaxes_kirki
EV_0068_Charaxes_northcotti
ET0210_Charaxes_viola
ABRI_012_Charaxes_mafuga
ABRI_016_Charaxes_turlini
EV_0052_Charaxes_berkeleyi
EV_0050_Charaxes_sidamo
EV_0058_Charaxes_galawadiwosi
EV_0059_Charaxes_howarthi
KAP071_Charaxes_virilis
KAP507_Charaxes_plantroui
ABRI_004_Charaxes_nichetes
NW118_11_Charaxes_porthos
EV_0062_Charaxes_mycerina
KAP228_Charaxes_zelica
KAP506_Charaxes_lycurgus
FM_15_Euxanthe_trajanus
EV_0064_Euxanthe_tiberius
EV_0066_Euxanthe_madagascariensis
NW131_10_Euxanthe_eurinome
EV_0065_Euxanthe_wakefieldi
NW103_15_Euxanthe_crossleyi
KAP165_Charaxes_zingha
EV_0039_Charaxes_hadrianus
KAP163_Charaxes_protoclea
KAP050_Charaxes_boueti
NW107_11_Charaxes_cynthia
EV_0020_Charaxes_lasti
KAP299_Charaxes_fulvescens
KAP503_Charaxes_varanes
ABRI_010_Charaxes_acuminatus
NW78_3_Charaxes_castor
EV_0022_Charaxes_jasius
EV_0023_Charaxes_legeri
KAP069_Charaxes_lucretius
KAP081_Charaxes_brutus
EV_0029_Charaxes_ansorgei
ABRI_019_Charaxes_eudoxus
EV_0035_Charaxes_richelmanni
KAP501_Charaxes_pollux
EV_0025_Charaxes_lactetinctus
EV_0034_Charaxes_ducarmei
ABRI_032_Charaxes_druceanus
EV_0006_Charaxes_fournierae
EV_0007_Charaxes_acraeoides
EV_0002_Charaxes_nobilis
EV_0001_Charaxes_superbus
EV_0038_Charaxes_imperialis
EV_0037_Charaxes_pythodoris
KAP509_Charaxes_numenes
KAP280_Charaxes_ameliae
ABRI_005_Charaxes_phenix
KAP290_Charaxes_bipunctatus
ABRI_031_Charaxes_bohemani
EV_0036_Charaxes_mixtus
KAP502_Charaxes_smaragdalis
KAP098_Charaxes_tiridates
EV_0032_Charaxes_cithaeron
EV_0033_Charaxes_xiphares
EV_0031_Charaxes_nandina
ABRI_020_Charaxes_cowani
EV_0018_Charaxes_antamboulou
KAP273_Charaxes_candiope
CJM_148_001_Charaxes_marki
CJM_171_001_Charaxes_ocellatus
CJM_173_001_Charaxes_orilus
CJM_194_001_Charaxes_harmodius
CJM_186_001_Charaxes_distanti
CJM_199_004_Charaxes_mars
CJM_191_007_Charaxes_nitebis
CJM_175_002_Charaxes_amycus
CJM_202_001_Charaxes_sangana
CJM_179_002_Charaxes_antonius
CJM_189_001_Charaxes_elwesi
CJM_196_001_Charaxes_eurialus
CJM_177_002_Charaxes_madensis
CJM_181_007_Charaxes_latona
CJM_166_001_Charaxes_setan
CJM_192_001_Charaxes_musashi
CJM_170_005_Charaxes_affinis
CJM_185_001_Charaxes_kahruba
CJM_183_007_Charaxes_repetitus
CJM_209_002_Charaxes_plateni
CJM_188_001_Charaxes_bupalus
CJM_205_003_Charaxes_aristogiton
CJM_195_003_Charaxes_borneensis
NW134_10_Charaxes_bernardus
UN0479_Charaxes_marmax
CJM_206_002_Charaxes_fervens
CJM_210_003_Charaxes_bajula
CJM_212_001_Charaxes_durnfordi
CJM_204_002_Charaxes_psaphon

**Figure SII. 4.** (page before). Maximum clade credibility tree for the Charaxinae subclade from BEAST. Posterior probabilities are indicated at the nodes.

e) Cyrestinae

*Dataset* – The dataset for the Cyrestinae consisted of 28 taxa to which two outgroups were added: *Antanartia delius* (Antanartia), *Chitoria ulupi* (Apaturinae).

*Partition Finder* – Partition Finder identified 11 subsets.

| Subset # | Substitution model | Gene fragments and codon positions |
| --- | --- | --- |
| Subset 1 | GTR+G | wingless_pos1, RpS5_pos1, ArgKin_pos1, RpS2_pos1, MDH_pos1, CAD_pos1 |
| Subset 2 | HKY+I | DDC_pos2, ArgKin_pos2, CAD_pos2, IDH_pos2 |
| Subset 3 | GTR+G | EF1a_pos3, ArgKin_pos3 |
| Subset 4 | HKY+I+G | MDH_pos3, CAD_pos3 |
| Subset 5 | GTR+I+G | COI-end_pos3, COI-begin_pos3 |
| Subset 6 | TRN+I+G | COI-end_pos1, COI-begin_pos1 |
| Subset 7 | HKY+I | COI-begin_pos2, COI-end_pos2, RpS5_pos2, GAPDH_pos2, MDH_pos2, EF1a_pos2 |
| Subset 8 | TRN+G | wingless_pos3, DDC_pos3 |
| Subset 9 | TRN+I+G | wingless_pos2, DDC_pos1, IDH_pos1, GAPDH_pos1, EF1a_pos1 |
| Subset 10 | GTR+G | RpS5_pos3, RpS2_pos3, IDH_pos3, GAPDH_pos3 |
| Subset 11 | JC | RpS2_pos2 |

*BEAST analysis* – In order to improve the quality of our runs we replaced the default priors for rates of substitutions by uniform prior ranging between 0 and 10 for the following cases: subset5. at, cg, gt. We used one molecular clock per subset identified by Partition Finder and obtained good mixing and convergence. We used a birth-death tree prior. We performed two runs of 60 million generations, sampling trees and parameters every 6000 generations.

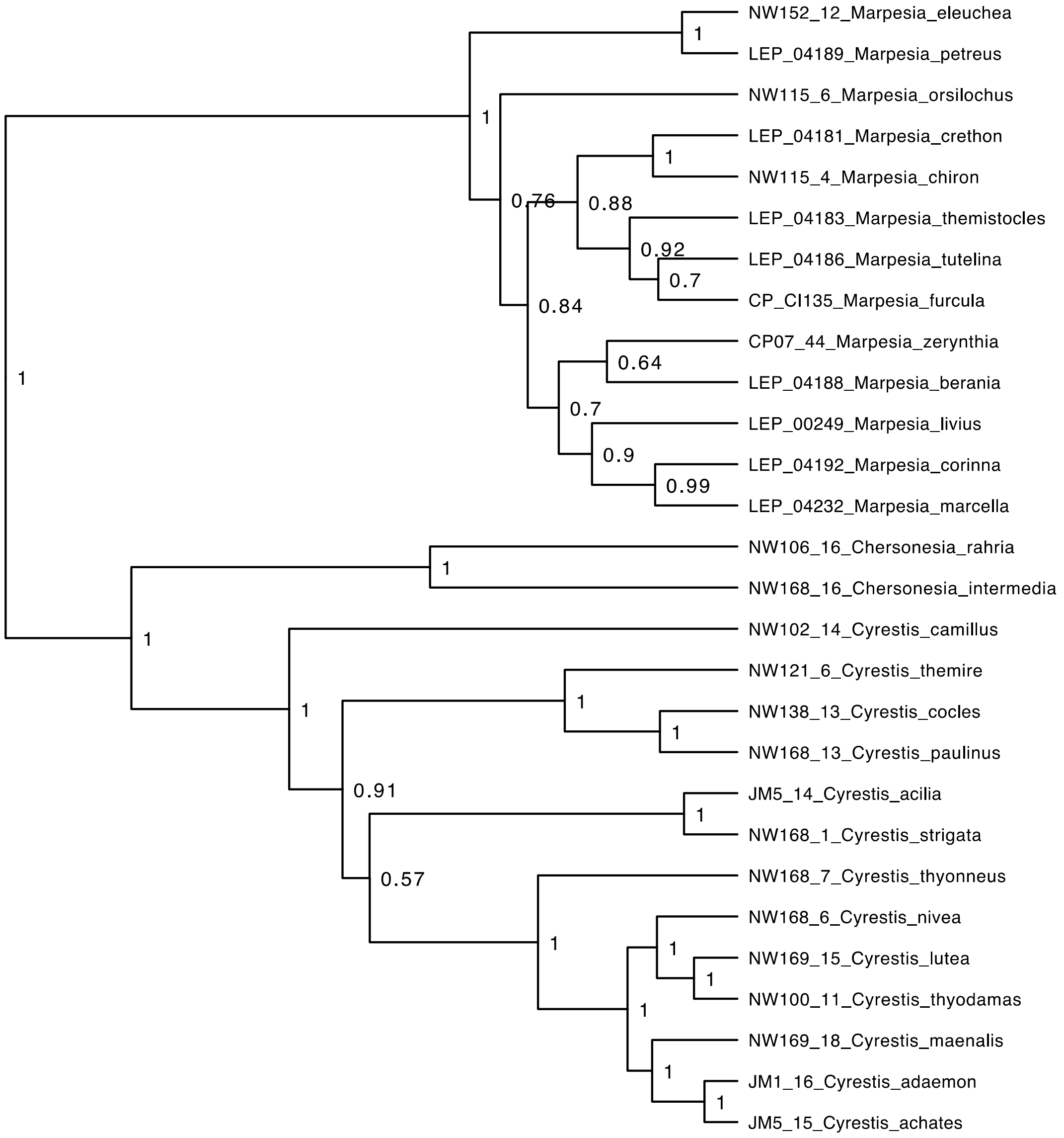


**Figure SII. 5.** Maximum clade credibility tree for the Cyrestinae subclade from BEAST. Posterior probabilities are indicated at the nodes.

f) Danaini

*Dataset* – The dataset for the Danaini consisted of 46 taxa to which two outgroups were added: *Tellervo zoilus* (Tellervini), *Libythea celtis* (Libytheinae).

*Partition Finder* – Partition Finder identified 11 subsets.

| Subset # | Substitution model | Gene fragments and codon positions |
|---|---|---|
| Subset 1 | GTR+G | wingless_pos1, RpS5_pos1, ArgKin_pos1, RpS2_pos1, MDH_pos1, CAD_pos1 |
| Subset 2 | HKY+I | DDC_pos2, ArgKin_pos2, IDH_pos2, CAD_pos2 |
| Subset 3 | GTR+G | EF1a_pos3, ArgKin_pos3 |
| Subset 4 | HKY+I+G | MDH_pos3, CAD_pos3 |

| | | |
|---|---|---|
| Subset 5 | GTR+I+G | COI-end_pos3, COI-begin_pos3 |
| Subset 6 | TRN+I+G | COI-end_pos1, COI-begin_pos1 |
| Subset 7 | HKY+I | COI-begin_pos2, COI-end_pos2, RpS5_pos2, GAPDH_pos2, MDH_pos2, EF1a_pos2 |
| Subset 8 | TRNEF+G | DDC_pos3, wingless_pos3 |
| Subset 9 | TRN+I+G | wingless_pos2, DDC_pos1, IDH_pos1, GAPDH_pos1, EF1a_pos1 |
| Subset 10 | GTR+G | RpS5_pos3, RpS2_pos3, IDH_pos3, GAPDH_pos3 |
| Subset 11 | JC | RpS2_pos2 |

*BEAST analysis* – In order to improve the quality of our runs we replaced the default priors for rates of substitutions by uniform prior ranging between 0 and 10 for the following cases: subset5.at, cg, gt, subset10.cg, subset1.cg. We used a single molecular clock linked across all partition to obtain good mixing and convergence. We used a birth-death tree prior. We performed two runs of 60 million generations, sampling trees and parameters every 7000 generations.

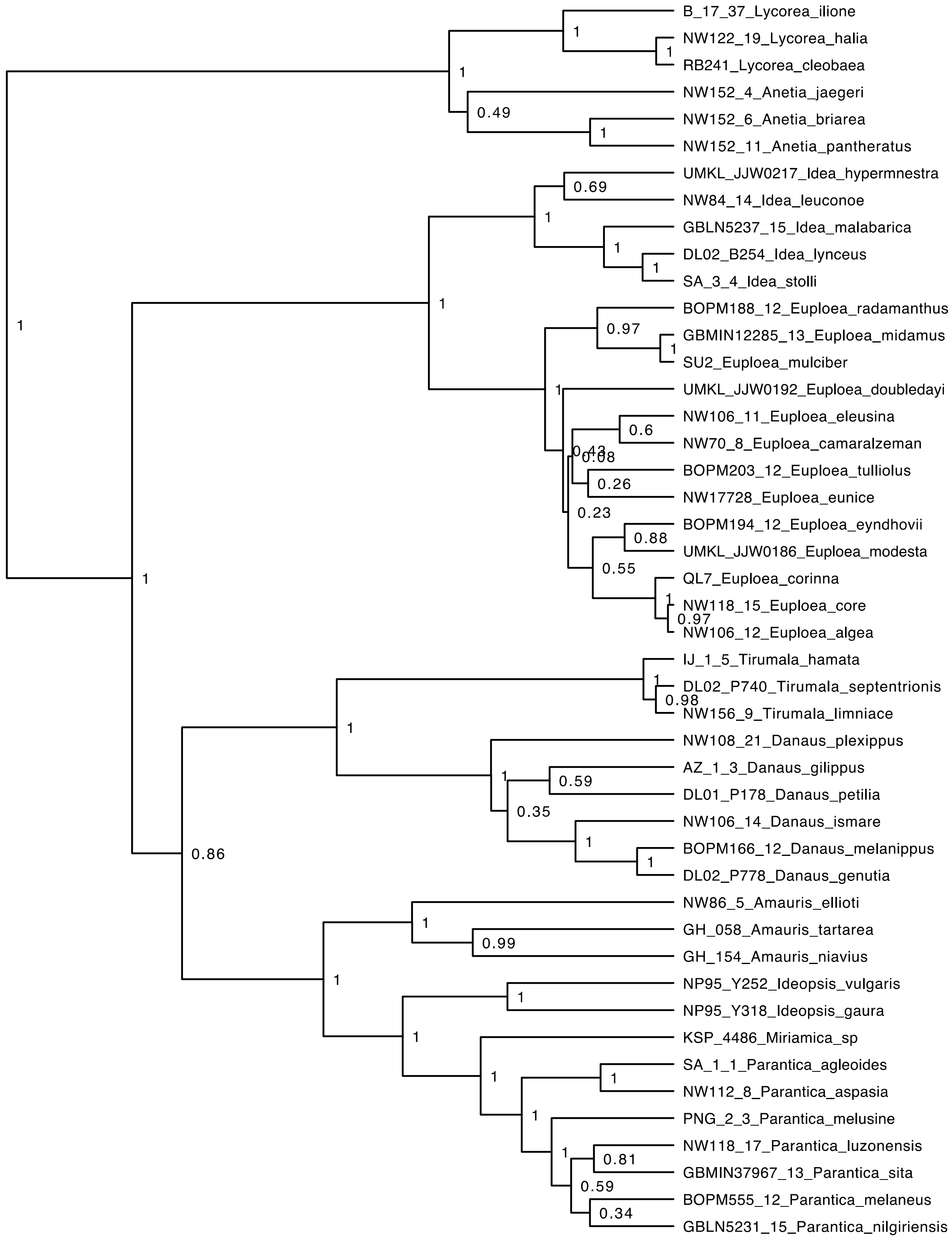


**Figure SII. 6.** Maximum clade credibility tree for the Danaini subclade from BEAST. Posterior probabilities are indicated at the nodes.

g) <u>Heliconiinae</u>

*Dataset* – The dataset for the Heliconiinae consisted of 248 taxa to which two outgroups were added: *Pseudergolis wedah* (Pseudergolinae), *Cymothoe caenis* (Limenitidinae).

*Partition Finder* – Partition Finder identified 13 subsets.

| Subset # | Substitution model | Gene fragments and codon positions |
|---|---|---|
| Subset 1 | TRN+I+G | GAPDH_pos1, EF1a_pos1, ArgKin_pos1, RpS2_pos1, RpS5_pos1 |
| Subset 2 | TRN+I+G | RpS2_pos2, RpS5_pos2, EF1a_pos2, GAPDH_pos2, ArgKin_pos2 |
| Subset 3 | GTR+G | ArgKin_pos3 |
| Subset 4 | HKY+I+G | MDH_pos3, CAD_pos3 |
| Subset 5 | GTR+I+G | CAD_pos1, MDH_pos1, IDH_pos1 |
| Subset 6 | GTR+I+G | DDC_pos2, IDH_pos2, CAD_pos2, MDH_pos2, COI-begin_pos2, COI-end_pos2 |
| Subset 7 | GTR+G | COI-begin_pos3 |
| Subset 8 | TRN+I+G | COI-end_pos1, COI-begin_pos1 |
| Subset 9 | GTR+G | COI-end_pos3 |
| Subset 10 | GTR+I+G | IDH_pos3, DDC_pos3 |
| Subset 11 | SYM+G | wingless_pos2, DDC_pos1, wingless_pos1 |
| Subset 12 | GTR+I+G | RpS2_pos3, wingless_pos3, EF1a_pos3, RpS5_pos3 |
| Subset 13 | TRN+I+G | GAPDH_pos3 |

*BEAST analysis* – In order to improve the quality of our runs we replaced the default priors for rates of substitutions by uniform prior ranging between 0 and 10 for the following cases: subset9.cg. We used one molecular clock per subset identified by Partition Finder and obtained good mixing and convergence. We used a birth-death tree prior. We performed three runs of 100 million generations, sampling trees and parameters every 10000 generations.

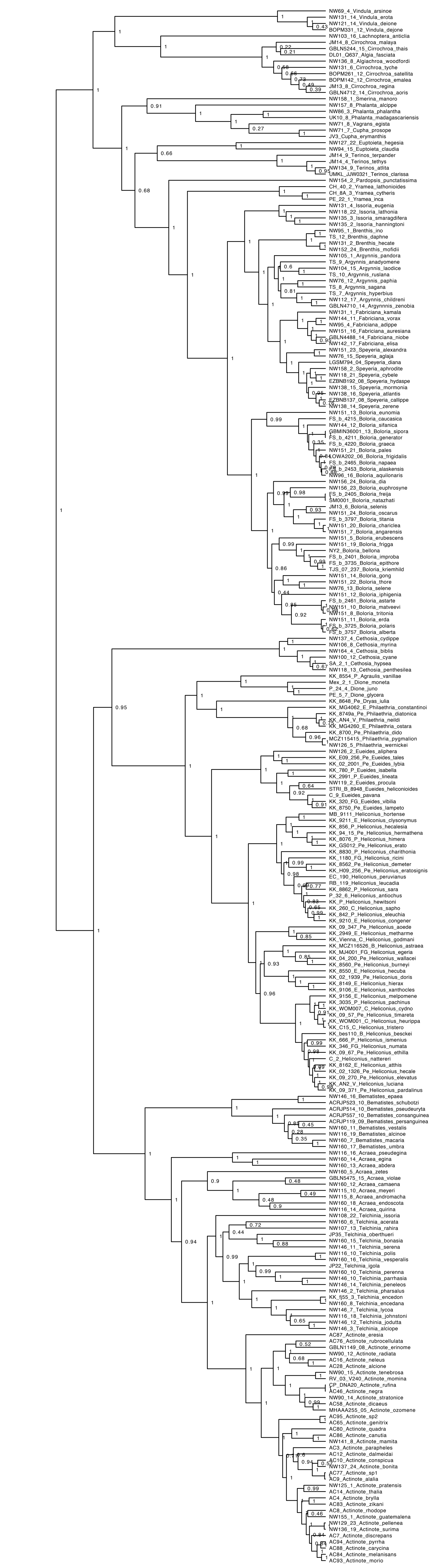
NW69_4_Vindula_arsinoe
NW131_14_Vindula_erota
NW121_14_Vindula_deione
BOPM331_12_Vindula_dejone
NW103_16_Lachnoptera_anticlia
JM14_8_Cirrochroa_malaya
GBLN5244_15_Cirrochroa_thais
DL01_Q637_Algia_fasciata
NW136_8_Algiachroa_woodfordi
NW131_6_Cirrochroa_tyche
BOPM261_12_Cirrochroa_satellita
BOPM142_12_Cirrochroa_emalea
JM13_8_Cirrochroa_regina
GBLN4712_14_Cirrochroa_aoris
NW158_1_Smerina_manoro
NW157_8_Phalanta_alcippe
NW86_3_Phalanta_phalantha
UK10_8_Phalanta_madagascariensis
NW71_8_Vagrans_egista
NW71_7_Cupha_prosope
JV3_Cupha_erymanthis
NW127_22_Euptoieta_hegesia
NW94_15_Euptoieta_claudia
JM14_9_Terinos_terpander
JM14_4_Terinos_tethys
NW134_9_Terinos_atlita
UMKL_JJW0321_Terinos_clarissa
NW154_2_Pardopsis_punctatissima
CH_40_2_Yramea_lathonioides
CH_8A_3_Yramea_cytheris
PE_22_1_Yramea_inca
NW131_4_Issoria_eugenia
NW118_22_Issoria_lathonia
NW135_3_Issoria_smaragdifera
NW135_2_Issoria_hanningtoni
NW95_1_Brenthis_ino
TS_12_Brenthis_daphne
NW131_2_Brenthis_hecate
NW152_24_Brenthis_mofidii
NW105_1_Argynnis_pandora
TS_9_Argynnis_anadyomene
NW104_15_Argynnis_laodice
TS_10_Argynnis_ruslana
NW76_12_Argynnis_paphia
TS_8_Argynnis_sagana
TS_7_Argynnis_hyperbius
NW112_17_Argynnis_childreni
GBLN4710_14_Argynnis_zenobia
NW131_1_Fabriciana_kamala
NW144_11_Fabriciana_vorax
NW95_4_Fabriciana_adippe
NW151_16_Fabriciana_auresiana
GBLN4488_14_Fabriciana_niobe
NW142_17_Fabriciana_elisa
NW151_23_Speyeria_alexandra
NW76_15_Speyeria_aglaja
LGSM794_04_Speyeria_diana
NW158_2_Speyeria_aphrodite
NW118_21_Speyeria_cybele
EZBNB192_08_Speyeria_hydaspe
NW138_15_Speyeria_mormonia
NW138_16_Speyeria_atlantis
EZBNB137_08_Speyeria_callippe
NW138_14_Speyeria_zerene
NW151_13_Boloria_eunomia
FS_b_4215_Boloria_caucasica
NW144_12_Boloria_sifanica
GBMIN36001_13_Boloria_sipora
FS_b_4211_Boloria_generator
FS_b_4220_Boloria_graeca
NW151_21_Boloria_pales
LOWA202_06_Boloria_frigidalis
FS_b_2465_Boloria_napaea
FS_b_2453_Boloria_alaskensis
NW96_16_Boloria_aquilonaris
NW156_24_Boloria_dia
NW156_23_Boloria_euphrosyne
FS_b_2405_Boloria_freija
SM0001_Boloria_natazhati
JM13_6_Boloria_selenis
NW151_24_Boloria_oscarus
FS_b_3797_Boloria_titania
NW151_20_Boloria_chariclea
NW151_7_Boloria_angarensis
NW151_5_Boloria_erubescens
NW151_19_Boloria_frigga
NY2_Boloria_bellona
FS_b_2401_Boloria_improba
FS_b_3735_Boloria_epithore
TJS_07_237_Boloria_kriemhild
NW151_14_Boloria_gong
NW151_22_Boloria_thore
NW76_13_Boloria_selene
NW151_12_Boloria_iphigenia
FS_b_2461_Boloria_astarte
NW151_10_Boloria_matveevi
NW151_8_Boloria_tritonia
NW151_11_Boloria_erda
FS_b_3725_Boloria_polaris
FS_b_3757_Boloria_alberta
NW137_4_Cethosia_cydippe
NW106_8_Cethosia_myrina
NW164_4_Cethosia_biblis
NW100_12_Cethosia_cyane
SA_2_1_Cethosia_hypsea
NW118_13_Cethosia_penthesilea
KK_8554_P_Agraulis_vanillae
Mex_2_1_Dione_moneta
P_24_4_Dione_juno
PE_5_7_Dione_glycera
KK_8648_Pe_Dryas_iulia
KK_MG4062_E_Philaethria_constantinoi
KK_8749a_Pe_Philaethria_diatonica
KK_AN4_V_Philaethria_neildi
KK_MG4260_E_Philaethria_ostara
KK_8700_Pe_Philaethria_dido
MCZ115415_Philaethria_pygmalion
NW126_5_Philaethria_wernickei
NW126_2_Eueides_aliphera
KK_E09_256_Pe_Eueides_tales
KK_02_2001_Pe_Eueides_lybia
KK_780_P_Eueides_isabella
KK_2991_P_Eueides_lineata
NW119_2_Eueides_procula
STRI_B_8948_Eueides_heliconioides
C_9_Eueides_pavana
KK_320_FG_Eueides_vibilia
KK_8750_Pe_Eueides_lampeto
MB_9111_Heliconius_hortense
KK_9211_E_Heliconius_clysonymus
KK_856_P_Heliconius_hecalesia
KK_94_15_Pe_Heliconius_hermathena
KK_8076_P_Heliconius_himera
KK_GS012_Pe_Heliconius_erato
KK_8830_P_Heliconius_charithonia
KK_1180_FG_Heliconius_ricini
KK_8562_Pe_Heliconius_demeter
KK_H09_256_Pe_Heliconius_eratosignis
EC_190_Heliconius_peruvianus
RB_119_Heliconius_leucadia
KK_8862_P_Heliconius_sara
P_32_6_Heliconius_antiochus
KK_P_Heliconius_hewitsoni
KK_260_C_Heliconius_sapho
KK_842_P_Heliconius_eleuchia
KK_9210_E_Heliconius_congener
KK_09_347_Pe_Heliconius_aoede
KK_2949_E_Heliconius_metharme
KK_Vienna_C_Heliconius_godmani
KK_MCZ116526_B_Heliconius_astraea
KK_MJ4001_FG_Heliconius_egeria
KK_04_200_Pe_Heliconius_wallacei
KK_8560_Pe_Heliconius_burneyi
KK_8550_E_Heliconius_hecuba
KK_02_1939_Pe_Heliconius_doris
KK_8149_E_Heliconius_hierax
KK_9106_E_Heliconius_xanthocles
KK_9156_E_Heliconius_melpomene
KK_3035_P_Heliconius_pachinus
KK_WOM007_C_Heliconius_cydno
KK_09_57_Pe_Heliconius_timareta
KK_WOM001_C_Heliconius_heurippa
KK_C15_C_Heliconius_tristero
KK_bes110_B_Heliconius_besckei
KK_666_P_Heliconius_ismenius
KK_346_FG_Heliconius_numata
KK_09_67_Pe_Heliconius_ethilla
C_2_Heliconius_nattereri
KK_8162_E_Heliconius_atthis
KK_02_1326_Pe_Heliconius_hecale
KK_09_270_Pe_Heliconius_elevatus
KK_AN2_V_Heliconius_luciana
KK_09_371_Pe_Heliconius_pardalinus
NW146_16_Bematistes_epaea
ACRJP523_10_Bematistes_schubotzi
ACRJP514_10_Bematistes_pseudeuryta
ACRJP557_10_Bematistes_consanguinea
ACRJP119_09_Bematistes_persanguinea
NW160_11_Bematistes_vestalis
NW116_19_Bematistes_alcinoe
NW160_7_Bematistes_macaria
NW160_17_Bematistes_umbra
NW116_16_Acraea_pseudegina
NW160_14_Acraea_egina
NW160_13_Acraea_abdera
NW160_5_Acraea_zetes
GBLN5475_15_Acraea_violae
NW160_12_Acraea_camaena
NW115_10_Acraea_meyeri
NW115_8_Acraea_andromacha
NW160_18_Acraea_endoscota
NW116_14_Acraea_quirina
NW108_22_Telchinia_issoria
NW160_6_Telchinia_acerata
NW107_13_Telchinia_rahira
JP35_Telchinia_oberthueri
NW160_15_Telchinia_bonasia
NW146_11_Telchinia_serena
NW116_10_Telchinia_polis
NW160_16_Telchinia_vesperalis
JP22_Telchinia_igola
NW160_10_Telchinia_perenna
NW146_10_Telchinia_parrhasia
NW146_14_Telchinia_peneleos
NW146_2_Telchinia_pharsalus
KK_fj55_3_Telchinia_encedon
NW160_8_Telchinia_encedana
NW146_7_Telchinia_lycoa
NW116_18_Telchinia_johnstoni
NW146_12_Telchinia_jodutta
NW146_3_Telchinia_alciope
AC87_Actinote_eresia
AC76_Actinote_rubrocellulata
GBLN1149_08_Actinote_erinome
NW90_12_Actinote_radiata
AC16_Actinote_neleus
AC28_Actinote_alcione
NW90_15_Actinote_tenebrosa
RV_03_V240_Actinote_momina
CP_DNA20_Actinote_rufina
AC46_Actinote_negra
NW90_14_Actinote_stratonice
AC58_Actinote_dicaeus
MHAAA255_05_Actinote_ozomene
AC95_Actinote_sp2
AC65_Actinote_genitrix
AC80_Actinote_quadra
AC86_Actinote_canutia
NW141_8_Actinote_mamita
AC3_Actinote_parapheles
AC12_Actinote_dalmeidai
AC10_Actinote_conspicua
NW137_24_Actinote_bonita
AC77_Actinote_sp1
AC9_Actinote_alalia
NW125_1_Actinote_pratensis
AC14_Actinote_thalia
AC4_Actinote_brylla
AC83_Actinote_zikani
AC8_Actinote_rhodope
NW155_1_Actinote_guatemalena
NW129_23_Actinote_pellenea
NW136_19_Actinote_surima
AC7_Actinote_discrepans
AC94_Actinote_pyrrha
AC88_Actinote_carycina
AC84_Actinote_melanisans
AC93_Actinote_morio

**Figure SII. 7.** (page before). Maximum clade credibility tree for the Heliconiinae subclade from BEAST. Posterior probabilities are indicated at the nodes.

h) Ithomiini

*Dataset* – The dataset for the Ithomiini consisted of 337 taxa to which two outgroups were added: *Danaus plexippus* (Danaini), *Tellervo zoilus* (Tellervini).

*Partition Finder* – Partition Finder identified 12 subsets.

| Subset # | Substitution model | Gene fragments and codon positions |
|---|---|---|
| Subset 1 | TRN+I+G | ArgKin_pos1, RpS2_pos1, EF1a_pos1 |
| Subset 2 | GTR+I+G | RpS5_pos2, GAPDH_pos2, DDC_pos2, CAD_pos2, IDH_pos2, ArgKin_pos2, EF1a_pos2 |
| Subset 3 | GTR | ArgKin_pos3 |
| Subset 4 | HKY+I+G | MDH_pos3, CAD_pos3 |
| Subset 5 | GTR+I+G | RpS5_pos1, GAPDH_pos1, wingless_pos1, MDH_pos1, CAD_pos1, IDH_pos1, DDC_pos1, wingless_pos2 |
| Subset 6 | GTR+I+G | COI-end_pos3, COI-begin_pos3 |
| Subset 7 | TRN+I+G | COI-end_pos1, COI-begin_pos1 |
| Subset 8 | TRN+I+G | MDH_pos2, COI-begin_pos2, COI-end_pos2 |
| Subset 9 | K80 | DDC_pos3 |
| Subset 10 | GTR+I+G | EF1a_pos3, wingless_pos3, RpS5_pos3 |
| Subset 11 | GTR+G | IDH_pos3, GAPDH_pos3, RpS2_pos3 |
| Subset 12 | JC+I | RpS2_pos2 |

*BEAST analysis* – In order to improve the quality of our runs we replaced the default priors for rates of substitutions by uniform prior ranging between 0 and 10 for the following cases: subset3.at, cg, gt, subset3.ag, cg. We used one molecular clock per subset identified by Partition Finder and obtained good mixing and convergence. We used a birth-death tree prior. We performed two runs of 100 million generations, sampling trees and parameters every 10000 generations.

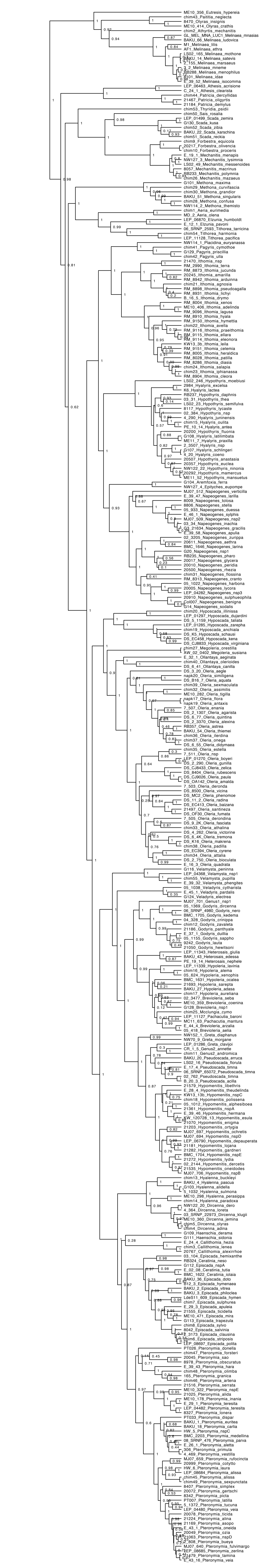

**Figure SII. 8.** Maximum clade credibility tree for the Ithomiini subclade from BEAST. Posterior probabilities are indicated at the nodes.

i) Libytheinae

*Dataset* – The dataset for the Libytheinae consisted of 10 taxa to which two outgroups were added: *Danaus plexippus* (Danaini), *Heliconius melpomene* (Heliconinae).

*Partition Finder* – Partition Finder identified 10 subsets.

| Subset # | Substitution model | Gene fragments and codon positions |
|---|---|---|
| Subset 1 | GTR+G | DDC_pos1, CAD_pos1, MDH_pos1, IDH_pos1, wingless_pos1, ArgKin_pos1, RpS5_pos1, RpS2_pos1, EF1a_pos1, GAPDH_pos1 |
| Subset 2 | HKY+I | RpS5_pos2, GAPDH_pos2, MDH_pos2, ArgKin_pos2, EF1a_pos2, IDH_pos2, DDC_pos2, CAD_pos2 |
| Subset 3 | HKY | ArgKin_pos3 |
| Subset 4 | HKY+G | IDH_pos3, MDH_pos3, CAD_pos3 |
| Subset 5 | HKY+G | COI-end_pos3, COI-begin_pos3 |
| Subset 6 | TRN+G | COI-end_pos1, COI-begin_pos1 |
| Subset 7 | HKY | COI-begin_pos2, COI-end_pos2 |
| Subset 8 | GTR | RpS2_pos3, DDC_pos3, GAPDH_pos3, RpS5_pos3, wingless_pos3 |
| Subset 9 | HKY+G | EF1a_pos3 |
| Subset 10 | JC | RpS2_pos2, wingless_pos2 |

*BEAST analysis* – In order to improve the quality of our runs we replaced the default priors for rates of substitutions by uniform prior ranging between 0 and 10 for the following cases: subset1.at, cg, gt. We used a single molecular clock linked across all partition to obtain good mixing and convergence. We used a birth-death tree prior. We performed two runs of 30 million generations, sampling trees and parameters every 3000 generations.

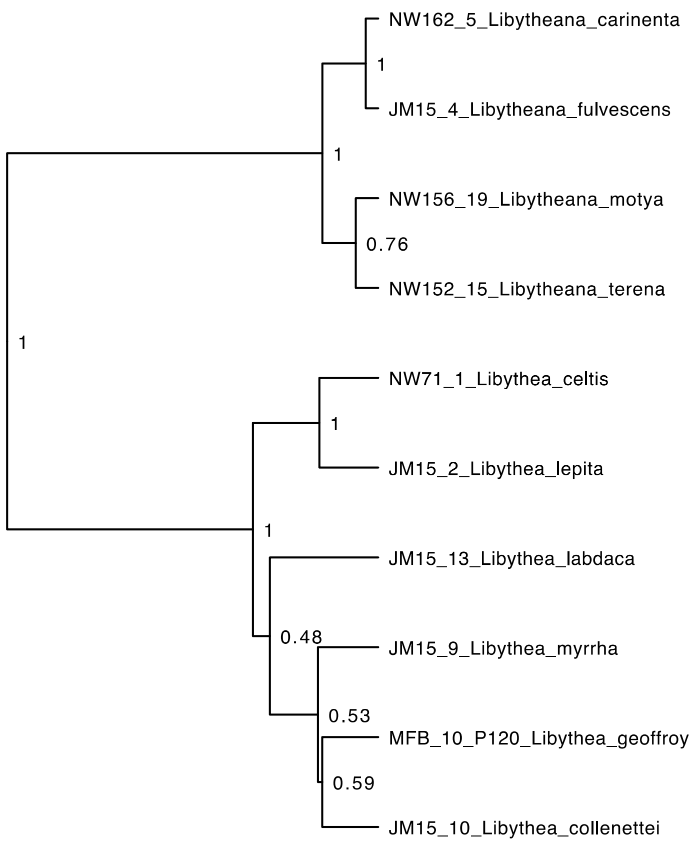


**Figure SII. 9.** Maximum clade credibility tree for the Libytheinae subclade from BEAST. Posterior probabilities are indicated at the nodes.

j) Limenitidinae

*Dataset* – The dataset for the Limenitidinae consisted of 311 taxa to which two outgroups were added: *Pseudergolis wedah* (Pseudergolinae), *Heliconius melpomene* (Heliconinae).

*Partition Finder* – Partition Finder identified 14 subsets.

| Subset # | Substitution model | Gene fragments and codon positions |
|---|---|---|
| Subset 1 | GTR+I+G | GAPDH_pos1, ArgKin_pos1, wingless_pos2, EF1a_pos1, RpS2_pos1 |
| Subset 2 | GTR+I+G | EF1a_pos2, ArgKin_pos2, DDC_pos2, RpS5_pos2, CAD_pos2, IDH_pos2, MDH_pos2, GAPDH_pos2 |
| Subset 3 | GTR+G | ArgKin_pos3 |
| Subset 4 | HKY+I+G | CAD_pos3, MDH_pos3 |
| Subset 5 | GTR+I+G | DDC_pos1, wingless_pos1, CAD_pos1, RpS5_pos1, MDH_pos1, IDH_pos1 |
| Subset 6 | GTR+I+G | COI-begin_pos3 |
| Subset 7 | GTR+I+G | COI-begin_pos1, COI-end_pos1 |
| Subset 8 | GTR+I+G | COI-begin_pos2, COI-end_pos2 |
| Subset 9 | GTR+G | COI-end_pos3 |
| Subset 10 | K80+I+G | DDC_pos3 |
| Subset 11 | GTR+I+G | EF1a_pos3 |
| Subset 12 | GTR+I+G | IDH_pos3, GAPDH_pos3 |

| | | |
|---|---|---|
| Subset 13 | JC | RpS2_pos2 |
| Subset 14 | GTR+I+G | RpS2_pos3, RpS5_pos3, wingless_pos3 |

*BEAST analysis* – After multiple attempts to modify priors we could not obtain a good convergence of the MCMC. Hence, we decided to fix the topology. We performed an analysis with IQtree to search for the best topology. The topology was then enforced in our BEAST analysis and we only estimated the relative branch lengths. We also replaced the default prior for rates of substitutions by uniform prior ranging between 0 and 10 for the following case: subset9.cg. We used two molecular clocks: one for the mitochondrial gene fragments and one for the nuclear gene fragments. We used a birth-death tree prior. We performed two runs of 50 million generations, sampling trees and parameters every 5000 generations.

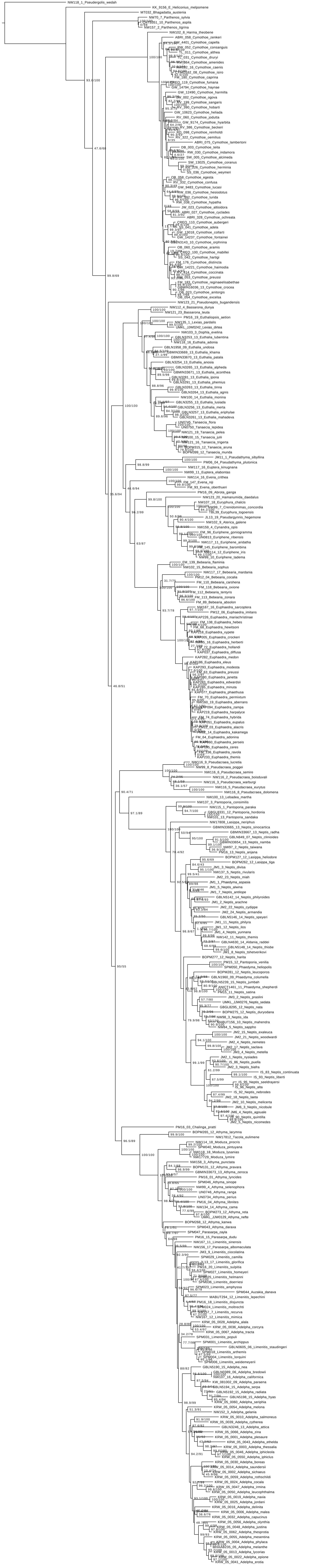

**Figure SII. 10.** (page before). Consensus tree for the Limenitidinae subclade from IQtree. Numbers at the nodes are SH-aLRT support (%) / ultrafast bootstrap support (%).

k) Satyrinae group II

*Dataset* – The dataset for the Satyrinae group II consisted of 169 taxa to which two outgroups were added: *Anaea troglodyta* (Charaxinae), *Brintesia circe* (Satyrinae).

*Partition Finder* – Partition Finder identified 10 subsets.

| Subset # | Substitution model | Gene fragments and codon positions |
|---|---|---|
| Subset 1 | TRN+I+G | wingless_pos2, ArgKin_pos1, RpS2_pos1, EF1a_pos1 |
| Subset 2 | TRN+I+G | RpS2_pos2, EF1a_pos2, RpS5_pos2, ArgKin_pos2, GAPDH_pos2, DDC_pos2 |
| Subset 3 | HKY+I+G | DDC_pos3, ArgKin_pos3, wingless_pos3 |
| Subset 4 | HKY+I+G | CAD_pos3, MDH_pos3 |
| Subset 5 | GTR+I+G | GAPDH_pos1, IDH_pos1, RpS5_pos1, MDH_pos1, CAD_pos1, DDC_pos1, wingless_pos1 |
| Subset 6 | GTR+I+G | MDH_pos2, COI-begin_pos2, COI-end_pos2, CAD_pos2, IDH_pos2 |
| Subset 7 | GTR+G | COI-begin_pos3, COI-end_pos3 |
| Subset 8 | GTR+I+G | COI-begin_pos1, COI-end_pos1 |
| Subset 9 | GTR+I+G | GAPDH_pos3, RpS5_pos3, EF1a_pos3 |
| Subset 10 | GTR+G | RpS2_pos3, IDH_pos3 |

*BEAST analysis* – In order to improve the quality of our runs we replaced the default priors for rates of substitutions by uniform prior ranging between 0 and 10 for the following cases: sub7.ag, at, gt, cg. We used one molecular clock per subset identified by Partition Finder and obtained good mixing and convergence. We used a birth-death tree prior. We performed three runs of 100 million generations, sampling trees and parameters every 10000 generations.

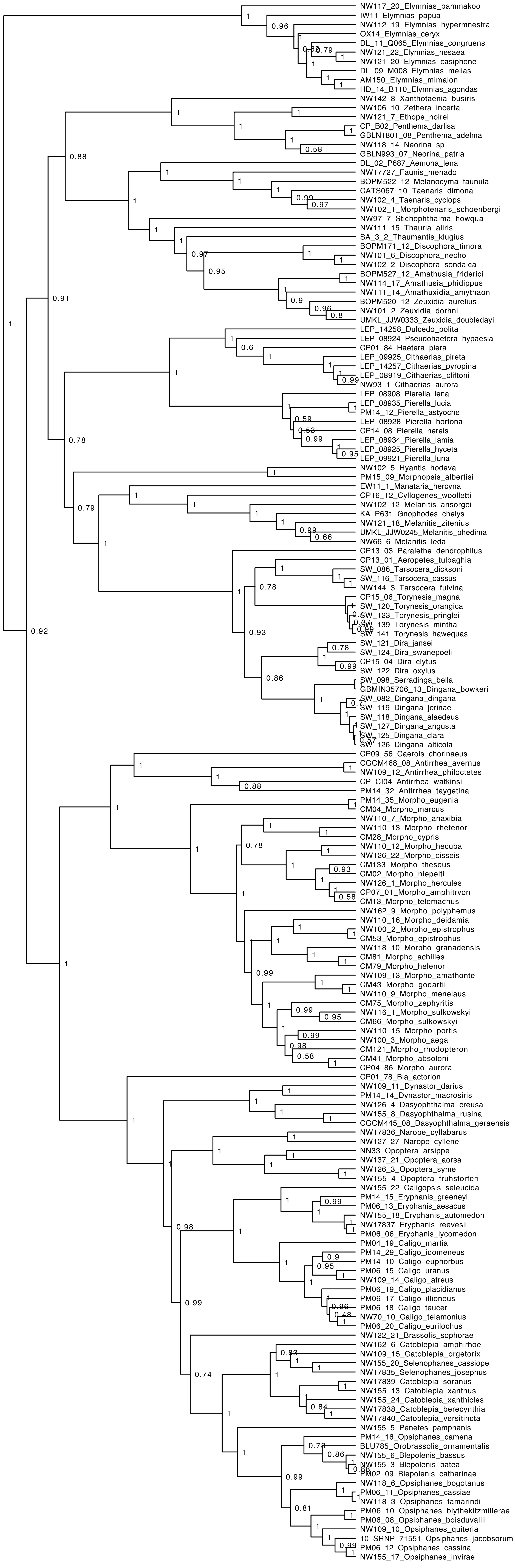
NW117_20_Elymnias_bammakoo
IW11_Elymnias_papua
NW112_19_Elymnias_hypermnestra
OX14_Elymnias_ceryx
DL_11_Q065_Elymnias_congruens
NW121_22_Elymnias_nesaea
NW121_20_Elymnias_casiphone
DL_09_M008_Elymnias_melias
AM150_Elymnias_mimalon
HD_14_B110_Elymnias_agondas
NW142_8_Xanthotaenia_busiris
NW106_10_Zethera_incerta
NW121_7_Ethope_noirei
CP_B02_Penthema_darlisa
GBLN1801_08_Penthema_adelma
NW118_14_Neorina_sp
GBLN993_07_Neorina_patria
DL_02_P687_Aemona_lena
NW17727_Faunis_menado
BOPM522_12_Melanocyma_faunula
CATS067_10_Taenaris_dimona
NW102_4_Taenaris_cyclops
NW102_1_Morphotenaris_schoenbergi
NW97_7_Stichophthalma_howqua
NW111_15_Thauria_aliris
SA_3_2_Thaumantis_klugius
BOPM171_12_Discophora_timora
NW101_6_Discophora_necho
NW102_2_Discophora_sondaica
BOPM527_12_Amathusia_friderici
NW114_17_Amathusia_phidippus
NW111_14_Amathuxidia_amythaon
BOPM520_12_Zeuxidia_aurelius
NW101_2_Zeuxidia_dorhni
UMKL_JJW0333_Zeuxidia_doubledayi
LEP_14258_Dulcedo_polita
LEP_08924_Pseudohaetera_hypaesia
CP01_84_Haetera_piera
LEP_09925_Cithaerias_pireta
LEP_14257_Cithaerias_pyropina
LEP_08919_Cithaerias_cliftoni
NW93_1_Cithaerias_aurora
LEP_08908_Pierella_lena
LEP_08935_Pierella_lucia
PM14_12_Pierella_astyoche
LEP_08928_Pierella_hortona
CP14_08_Pierella_nereis
LEP_08934_Pierella_lamia
LEP_08925_Pierella_hyceta
LEP_09921_Pierella_luna
NW102_5_Hyantis_hodeva
PM15_09_Morphopsis_albertisi
EW11_1_Manataria_hercyna
CP16_12_Cyllogenes_woolletti
NW102_12_Melanitis_ansorgei
KA_P631_Gnophodes_chelys
NW121_18_Melanitis_zitenius
UMKL_JJW0245_Melanitis_phedima
NW66_6_Melanitis_leda
CP13_03_Paralethe_dendrophilus
CP13_01_Aeropetes_tulbaghia
SW_086_Tarsocera_dicksoni
SW_116_Tarsocera_cassus
NW144_3_Tarsocera_fulvina
CP15_06_Torynesis_magna
SW_120_Torynesis_orangica
SW_123_Torynesis_pringlei
SW_139_Torynesis_mintha
SW_141_Torynesis_hawequas
SW_121_Dira_jansei
SW_124_Dira_swanepoeli
CP15_04_Dira_clytus
SW_122_Dira_oxylus
SW_098_Serradinga_bella
GBMIN35706_13_Dingana_bowkeri
SW_082_Dingana_dingana
SW_119_Dingana_jerinae
SW_118_Dingana_alaedeus
SW_127_Dingana_angusta
SW_125_Dingana_clara
SW_126_Dingana_alticola
CP09_56_Caerois_chorinaeus
CGCM468_08_Antirrhea_avernus
NW109_12_Antirrhea_philoctetes
CP_CI04_Antirrhea_watkinsi
PM14_32_Antirrhea_taygetina
PM14_35_Morpho_eugenia
CM04_Morpho_marcus
NW110_7_Morpho_anaxibia
NW110_13_Morpho_rhetenor
CM28_Morpho_cypris
NW110_12_Morpho_hecuba
NW126_22_Morpho_cisseis
CM133_Morpho_theseus
CM02_Morpho_niepelti
NW126_1_Morpho_hercules
CP07_01_Morpho_amphitryon
CM13_Morpho_telemachus
NW162_9_Morpho_polyphemus
NW110_16_Morpho_deidamia
NW100_2_Morpho_epistrophus
CM53_Morpho_epistrophus
NW118_10_Morpho_granadensis
CM81_Morpho_achilles
CM79_Morpho_helenor
NW109_13_Morpho_amathonte
CM43_Morpho_godartii
NW110_9_Morpho_menelaus
CM75_Morpho_zephyritis
NW116_1_Morpho_sulkowskyi
CM66_Morpho_sulkowskyi
NW110_15_Morpho_portis
NW100_3_Morpho_aega
CM121_Morpho_rhodopteron
CM41_Morpho_absoloni
CP04_86_Morpho_aurora
CP01_78_Bia_actorion
NW109_11_Dynastor_darius
PM14_14_Dynastor_macrosiris
NW126_4_Dasyophthalma_creusa
NW155_8_Dasyophthalma_rusina
CGCM445_08_Dasyophthalma_geraensis
NW17836_Narope_cyllabarus
NW127_27_Narope_cyllene
NN33_Opoptera_arsippe
NW137_21_Opoptera_aorsa
NW126_3_Opoptera_syme
NW155_4_Opoptera_fruhstorferi
NW155_22_Caligopsis_seleucida
PM14_15_Eryphanis_greeneyi
PM06_13_Eryphanis_aesacus
NW155_18_Eryphanis_automedon
NW17837_Eryphanis_reevesii
PM06_06_Eryphanis_lycomedon
PM04_19_Caligo_martia
PM14_29_Caligo_idomeneus
PM14_10_Caligo_euphorbus
PM06_15_Caligo_uranus
NW109_14_Caligo_atreus
PM06_19_Caligo_placidianus
PM06_17_Caligo_illioneus
PM06_18_Caligo_teucer
NW70_10_Caligo_telamonius
PM06_20_Caligo_eurilochus
NW122_21_Brassolis_sophorae
NW162_6_Catoblepia_amphirhoe
NW109_15_Catoblepia_orgetorix
NW155_20_Selenophanes_cassiope
NW17835_Selenophanes_josephus
NW17839_Catoblepia_soranus
NW155_13_Catoblepia_xanthus
NW155_24_Catoblepia_xanthicles
NW17838_Catoblepia_berecynthia
NW17840_Catoblepia_versitincta
NW155_5_Penetes_pamphanis
PM14_16_Opsiphanes_camena
BLU785_Orobrassolis_ornamentalis
NW155_6_Blepolenis_bassus
NW155_3_Blepolenis_batea
PM02_09_Blepolenis_catharinae
NW118_6_Opsiphanes_bogotanus
PM06_11_Opsiphanes_cassiae
NW118_3_Opsiphanes_tamarindi
PM06_10_Opsiphanes_blythekitzmillerae
PM06_08_Opsiphanes_boisduvallii
NW109_10_Opsiphanes_quiteria
10_SRNP_71551_Opsiphanes_jacobsorum
PM06_12_Opsiphanes_cassina
NW155_17_Opsiphanes_invirae

**Figure SII. 11.** (page before). Maximum clade credibility tree for the Satyrinae group I subclade from BEAST. Posterior probabilities are indicated at the nodes.

l) Satyrinae group II

*Dataset* – The dataset for the Satyrinae group I consisted of 368 taxa to which two outgroups were added: *Amathusia phidippus* (Satyrinae), *Brintesia circe* (Satyrinae).

*Partition Finder* – Partition Finder identified 14 subsets.

| Subset # | Substitution model | Gene fragments and codon positions |
|---|---|---|
| Subset 1 | GTR+I+G | CAD_pos1, RpS5_pos1, MDH_pos1, IDH_pos1, GAPDH_pos1, ArgKin_pos1 |
| Subset 2 | HKY+I+G | wingless_pos2, ArgKin_pos2, EF1a_pos2, DDC_pos2 |
| Subset 3 | GTR+I+G | ArgKin_pos3, wingless_pos3 |
| Subset 4 | HKY+I+G | GAPDH_pos3, CAD_pos3, MDH_pos3 |
| Subset 5 | GTR+I+G | COI-begin_pos2, MDH_pos2, COI-end_pos2, CAD_pos2, IDH_pos2 |
| Subset 6 | GTR+I+G | COI-begin_pos3 |
| Subset 7 | GTR+I+G | COI-begin_pos1, COI-end_pos1 |
| Subset 8 | GTR+G | COI-end_pos3 |
| Subset 9 | GTR+I+G | DDC_pos3, IDH_pos3, RpS2_pos3, RpS5_pos3 |
| Subset 10 | SYM+I+G | RpS2_pos1, DDC_pos1 |
| Subset 11 | HKY+I+G | EF1a_pos3 |
| Subset 12 | TRN+I+G | wingless_pos1, EF1a_pos1 |
| Subset 13 | TRN+I+G | GAPDH_pos2, RpS5_pos2 |
| Subset 14 | SYM+I+G | RpS2_pos2 |

*BEAST analysis* – In order to improve the quality of our runs we replaced the default priors for rates of substitutions by uniform prior ranging between 0 and 10 for the following cases: sub14.at, gt and sub8.cg, gt. We used a single molecular clock linked across all partition to obtain good mixing and convergence. We used a birth-death tree prior. We performed two runs of 60 million generations, sampling trees and parameters every 6000 generations.

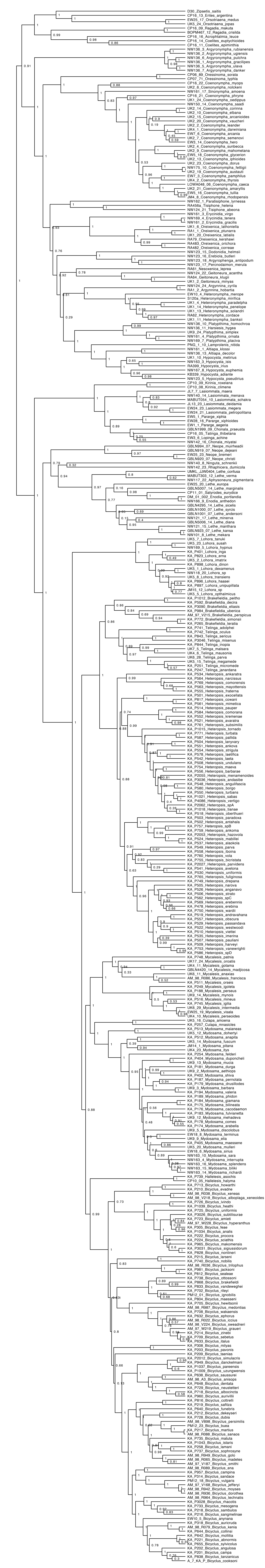

**Figure SII. 12.** (page before). Maximum clade credibility tree for the Satyrinae group II subclade from BEAST. Posterior probabilities are indicated at the nodes.

m) Nymphalinae

*Dataset* – The dataset for the Nymphalinae consisted of 359 taxa to which two outgroups were added: *Chitoria ulupi* (Apaturinae), *Marpesia eleuchea* (Cyrestinae).

*Partition Finder* – Partition Finder identified 13 subsets.

| Subset # | Substitution model | Gene fragments and codon positions |
|---|---|---|
| Subset 1 | GTR+I+G | GAPDH_pos1, IDH_pos1, ArgKin_pos1, wingless_pos1, MDH_pos1, CAD_pos1 |
| Subset 2 | TRN+I+G | RpS5_pos2, MDH_pos2, CAD_pos2, DDC_pos2, ArgKin_pos2, IDH_pos2 |
| Subset 3 | GTR+I+G | ArgKin_pos3, DDC_pos3 |
| Subset 4 | HKY+I+G | CAD_pos3, MDH_pos3 |
| Subset 5 | GTR+I+G | COI-begin_pos3, COI-end_pos3 |
| Subset 6 | TRN+I+G | COI-begin_pos1 |
| Subset 7 | GTR+I+G | COI-begin_pos2, COI-end_pos2 |
| Subset 8 | TRN+I+G | COI-end_pos1 |
| Subset 9 | K80+I+G | wingless_pos2, DDC_pos1 |
| Subset 10 | GTR+I+G | wingless_pos3, EF1a_pos3 |
| Subset 11 | TRN+I+G | RpS5_pos1, EF1a_pos1, RpS2_pos1 |
| Subset 12 | JC+I+G | EF1a_pos2, GAPDH_pos2, RpS2_pos2 |
| Subset 13 | GTR+I+G | RpS5_pos3, IDH_pos3, RpS2_pos3, GAPDH_pos3 |

*BEAST analysis* – After multiple attempts to modify priors, MCMC runs did not converge. Hence, we decided to fix the topology. We performed an analysis with IQtree to search for the best topology. The topology was then enforced in our BEAST analysis and we only estimated the relative branch lengths. We used one molecular clock per subset identified by Partition Finder and obtained good mixing and convergence. We used a birth-death tree prior. We performed two runs of 60 million generations, sampling trees and parameters every 60000 generations.

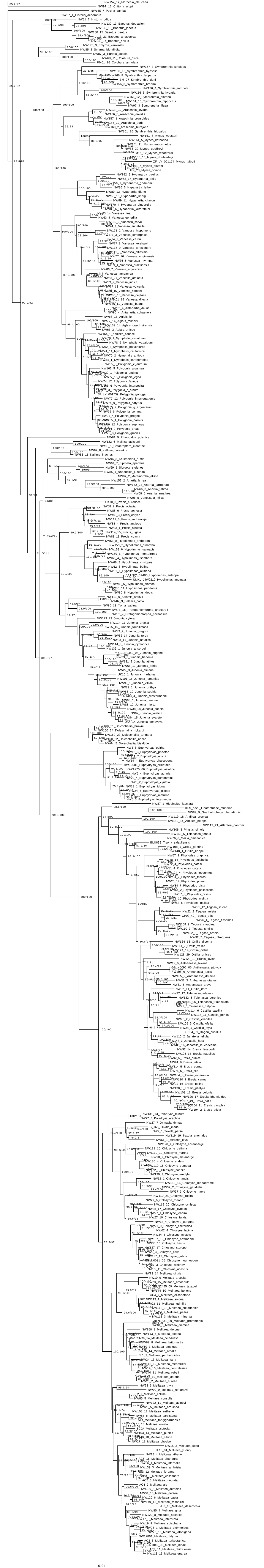
0.04

**Figure SII. 13.** (page before). Consensus tree for the Nymphalinae subclade from IQtree. Numbers at the nodes are SH-aLRT support (%) / ultrafast bootstrap support (%).

n) Pseudergolinae

*Dataset* – The dataset for the Pseudergolinae consisted of 6 taxa to which two outgroups were added: *Heliconius melpomene* (Satyrinae), *Marpesia eleuchea* (Cyrestinae).

*Partition Finder* – Partition Finder identified 10 subsets.

| Subset # | Substitution model | Gene fragments and codon positions |
|---|---|---|
| Subset 1 | TRN+G | wingless_pos1, DDC_pos1, ArgKin_pos1, MDH_pos1, RpS5_pos1, IDH_pos1, GAPDH_pos1, EF1a_pos1, RpS2_pos1 |
| Subset 2 | HKY+I | DDC_pos2, CAD_pos2, IDH_pos2, RpS5_pos2, ArgKin_pos2, EF1a_pos2, MDH_pos2, GAPDH_pos2 |
| Subset 3 | GTR+G | wingless_pos3, ArgKin_pos3, EF1a_pos3 |
| Subset 4 | TRN+G | CAD_pos3 |
| Subset 5 | GTR | CAD_pos1 |
| Subset 6 | HKY+G | COI-begin_pos3, COI-end_pos3 |
| Subset 7 | TRN+I | COI-begin_pos1, COI-end_pos1 |
| Subset 8 | HKY | COI-begin_pos2, COI-end_pos2 |
| Subset 9 | GTR+G | RpS5_pos3, DDC_pos3, RpS2_pos3, GAPDH_pos3, MDH_pos3, IDH_pos3 |
| Subset 10 | JC | RpS2_pos2, wingless_pos2 |

*BEAST analysis* – In order to improve the quality of our runs, we replaced the default priors for rates of substitutions by uniform prior ranging between 0 and 10 for the following cases: sub3.cg, sub5.at, cg, gt and sub9.ac, cg, gt. We used a single molecular clock linked across all partitions to obtain good mixing and convergence. We used a birth-death tree prior. We performed two runs of 30 million generations, sampling trees and parameters every 3000 generations.

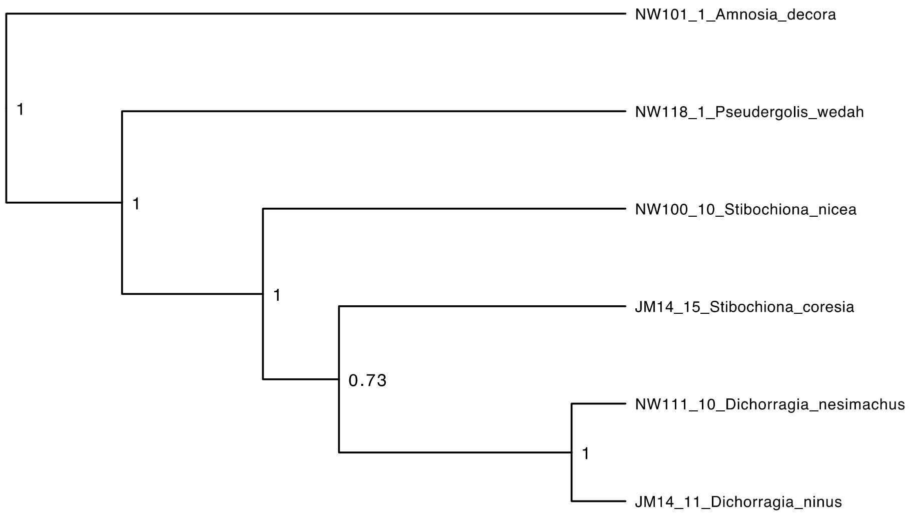


**Figure SII. 14.** Maximum clade credibility tree for the Pseudergolinae subclade from BEAST. Posterior probabilities are indicated at the nodes.

o) <u>Satyrini</u>

*Dataset* – The dataset for the Satyrini consisted of 560 taxa to which two outgroups were added: *Amathusia phidippus* (Satyrinae), *Bicyclus anynana* (Satyrinae).

*Partition Finder* – Partition Finder identified 17 subsets.

| Subset # | Substitution model | Gene fragments and codon positions |
|---|---|---|
| Subset 1 | TRN+I+G | ArgKin_pos1, RpS2_pos1, EF1a_pos1, MDH_pos1 |
| Subset 2 | HKY+I+G | ArgKin_pos2, RpS5_pos2, MDH_pos2, GAPDH_pos2, IDH_pos2 |
| Subset 3 | GTR+I+G | ArgKin_pos3, wingless_pos3 |
| Subset 4 | HKY+I+G | CAD_pos3, MDH_pos3 |
| Subset 5 | GTR+I+G | GAPDH_pos1, RpS5_pos1, CAD_pos1, IDH_pos1 |
| Subset 6 | HKY+I+G | DDC_pos2, CAD_pos2, COI-end_pos2 |
| Subset 7 | GTR+I+G | COI-begin_pos3 |
| Subset 8 | SYM+I | COI-begin_pos1 |
| Subset 9 | GTR+I+G | COI-begin_pos2 |
| Subset 10 | GTR+G | COI-end_pos3 |
| Subset 11 | HKY+I+G | COI-end_pos1 |
| Subset 12 | K80+G | DDC_pos3, RpS2_pos3 |
| Subset 13 | SYM+G | DDC_pos1, wingless_pos1 |
| Subset 14 | GTR+I+G | EF1a_pos3 |
| Subset 15 | K80+I+G | RpS2_pos2, wingless_pos2, EF1a_pos2 |
| Subset 16 | GTR+I+G | GAPDH_pos3 |
| Subset 17 | GTR+I+G | IDH_pos3, RpS5_pos3 |

*BEAST analysis* – We used a single molecular clock linked across all partitions to obtain good mixing and convergence. We used one molecular clock per subset identified by Partition

Finder and obtained good mixing and convergence. We used a birth-death tree prior. We performed two runs of 100 million generations, sampling trees and parameters every 10000 generations.

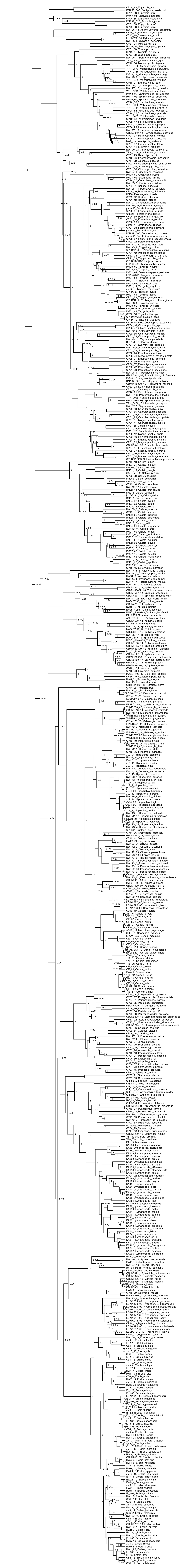

**Figure SII. 16.** (page before). Maximum clade credibility tree for the Satyrini subclade from BEAST. Posterior probabilities are indicated at the nodes.

#### *5 - Grafting procedure*

We built a posterior distribution of grafted trees by randomly sampling trees from the posterior distributions of both the backbone and the subclades. For each grafted tree, we (1) randomly picked a backbone from the posterior distribution without replacement, (2) randomly picked a tree from the posterior distribution of each subclade without replacement, (3) removed the two outgroups in the subclade tree, (4) rescaled the subclade trees to the crown age of the subclades in the backbone, (5) removed all the taxa representing the subclades in the backbone but one, (6) rescaled the branch length of the remaining taxa to the length of the stem leading to the subclades, and (7) grafted the subclade trees onto the corresponding stem branches. We repeated this process for 1000 random trees. Finally, we used TreeAnnotator v. 1.8.3 (Drummond et al 2012) to summarize the tree topology with median node age and 95% credibility interval of each node. The outgroups were finally removed and the resulting tree was used for all subsequent analyses.

### III - Estimation of speciation and extinction rate

We estimated the temporal dynamics of speciation and extinction rates across our phylogeny using BAMM 2.5 (Rabosky et al. 2013; Rabosky 2014; Rabosky et al. 2017). We accounted for missing species by specifying the sampling fraction at the genus level. We used the R-package BAMMtools (Rabosky et al. 2014) to generate initial priors for speciation and extinction rates. We set the expected number of rate shifts (Poisson rate parameter) to 0.4 for our main analysis. Given the concerns about the sensitivity of BAMM to priors on posteriors (Moore et al. 2016, but see Rabosky et al. 2017), we performed two additional BAMM analyses by setting the Poisson rate parameter to 0.2 (allowing fewer shifts) or 0.6 (allowing more shifts). All BAMM analyses ran 50 million generations with four reversible-jump MCMC, sampling the parameters every 50 000 generations. The outputs were then analyzed using the R-package BAMMtools. We checked that the MCMC converged with effective sample size after we discarded the first 10% of samples as burn-in. We extracted the best shift configuration by setting the expected number of shifts to five.

**Figure SIII. 1**. (see external figures) Results from BAMM analysis. Branches are colored according to the mean net diversification rate across the posterior distribution.

### IV - Inference of biogeographic history

We performed a maximum-likelihood estimate of geographical range evolution using the Dispersal-Extinction-Cladogenesis (DEC) model (Ree & Smith 2008) as implemented in the C++ version (Smith 2009) and recently extended to handle large phylogenetic trees such as the Nymphalidae, be faster even with a large number of possible ranges, and include new features (DECX model, Beeravolu & Condamine 2016). We ran the classical DEC model to estimate ancestral states at all nodes and the main dispersal events between the biogeographic regions. We designed a biogeographic model spanning the evolutionary history of Nymphalidae (Late Cretaceous). We assigned extant species to nine biogeographic regions: Western Nearctic, Eastern Nearctic, Western Palearctic, Eastern Palearctic, Neotropics, Afrotropics, India, Southeast Asia, and Australasia. We designed a time-stratified model in which both the adjacency matrices and dispersal matrices varied between time periods. Time was divided into five time periods: 100-80, 80-60, 60-30, 30-10, 10-0 million years to account for increasing or decreasing connectivity between biogeographic regions through time (Table SIV. 1. and Table SIV. 2.). We used the results of the full model (i.e., including both connectivity matrix and dispersal multipliers) for subsequent analyses, but we performed additional analyses with different models to test the effect of our choices on the ancestral-state estimation.

**Table SIV. 1.** Dispersal multipliers matrices for five time-intervals used for ancestral biogeographic state estimation using DECX. WN=Western Nearctic, EN=Eastern Nearctic, NT=Neotropic, WP=Western Palearctic, EN=Eastern Palearctic, AF=Afrotropic, MD=Madagascar, IN=India, SEA=South East Asia, AU=Australasia.

| **0 - 10 Myr** | WN | EN | NT | WP | EP | AF | MD | IN | SEA | AU |
|---|---|---|---|---|---|---|---|---|---|---|
| WN | 1 | 1 | 1 | 0.01 | 1 | 0 | 0 | 0 | 0.01 | 0.01 |
| EN | 1 | 1 | 1 | 0.1 | 0.01 | 0.01 | 0 | 0 | 0 | 0 |
| NT | 1 | 1 | 1 | 0.01 | 0.01 | 0.01 | 0 | 0 | 0 | 0.01 |
| WP | 0.01 | 0.1 | 0.01 | 1 | 1 | 1 | 0 | 1 | 0 | 0 |
| EP | 1 | 0.01 | 0.01 | 1 | 1 | 0.5 | 0 | 1 | 1 | 0.01 |
| AF | 0 | 0.01 | 0.01 | 1 | 0.5 | 1 | 1 | 0.5 | 0.5 | 0.01 |
| MD | 0 | 0 | 0 | 0 | 0 | 1 | 1 | 0.1 | 0.01 | 0.01 |
| IN | 0 | 0 | 0 | 1 | 1 | 0.5 | 0.1 | 1 | 1 | 0.01 |
| SEA | 0.01 | 0 | 0 | 0 | 1 | 0.5 | 0.01 | 1 | 1 | 1 |
| AU | 0.01 | 0 | 0.01 | 0 | 0.01 | 0.01 | 0.01 | 0.01 | 0.01 | 1 |

| **10 - 30 Myr** | WN | EN | NT | WP | EP | AF | MD | IN | SEA | AU |
|---|---|---|---|---|---|---|---|---|---|---|
| WN | 1 | 1 | 0.5 | 0.01 | 1 | 0 | 0 | 0 | 0.01 | 0.01 |
| EN | 1 | 1 | 0.5 | 0.1 | 0.01 | 0.01 | 0 | 0 | 0 | 0 |
| NT | 0.5 | 0.5 | 1 | 0.01 | 0.01 | 0.01 | 0 | 0 | 0 | 0.01 |
| WP | 0.01 | 0.1 | 0.01 | 1 | 1 | 0 | 0.1 | 0 | 0.01 | 0 |

| | | | | | | | | | | |
|---|---|---|---|---|---|---|---|---|---|---|
| EP | 1 | 0.01 | 0.01 | 1 | 1 | 0.5 | 0 | 0.5 | 1 | 0.01 |
| AF | 0 | 0.01 | 0.01 | 0 | 0.5 | 1 | 1 | 0.5 | 0.5 | 0.01 |
| MD | 0 | 0 | 0 | 0.1 | 0 | 1 | 1 | 0.1 | 0.01 | 0.01 |
| IN | 0 | 0 | 0 | 0 | 0.5 | 0.5 | 0.1 | 1 | 0.5 | 0.01 |
| SEA | 0.01 | 0 | 0 | 0.01 | 1 | 0.5 | 0.01 | 0.5 | 1 | 0.5 |
| AU | 0.01 | 0 | 0.01 | 0 | 0.01 | 0.01 | 0.01 | 0.01 | 0.5 | 1 |

| **30 - 60 Myr** | WN | EN | NT | WP | EP | AF | MD | IN | SEA | AU |
|---|---|---|---|---|---|---|---|---|---|---|
| WN | 1 | 1 | 0.1 | 0.01 | 1 | 0 | 0 | 0 | 0.01 | 0.01 |
| EN | 1 | 1 | 0.1 | 0.5 | 0.01 | 0.01 | 0 | 0 | 0 | 0 |
| NT | 0.1 | 0.1 | 1 | 0.01 | 0.01 | 0.1 | 0 | 0 | 0.01 | 0.1 |
| WP | 0.01 | 0.5 | 0.01 | 1 | 0.5 | 1 | 0 | 0.1 | 0 | 0 |
| EP | 1 | 0.01 | 0.01 | 0.5 | 1 | 0.1 | 0 | 0.1 | 1 | 0.01 |
| AF | 0 | 0.01 | 0.1 | 1 | 0.1 | 1 | 1 | 0.1 | 0.1 | 0.01 |
| MD | 0 | 0 | 0 | 0 | 0 | 1 | 1 | 0.1 | 0.01 | 0.01 |
| IN | 0 | 0 | 0 | 0.1 | 0.1 | 0.1 | 0.1 | 1 | 0.1 | 0.01 |
| SEA | 0.01 | 0 | 0.01 | 0 | 1 | 0.1 | 0.01 | 0.1 | 1 | 0.1 |
| AU | 0.01 | 0 | 0.1 | 0 | 0.01 | 0.01 | 0.01 | 0.01 | 0.1 | 1 |

| **60 - 80 Myr** | WN | EN | NT | WP | EP | AF | MD | IN | SEA | AU |
|---|---|---|---|---|---|---|---|---|---|---|
| WN | 1 | 1 | 0.1 | 0.01 | 1 | 0 | 0 | 0 | 0.01 | 0.01 |
| EN | 1 | 1 | 0.1 | 0.5 | 0.01 | 0.01 | 0 | 0 | 0 | 0 |
| NT | 0.1 | 0.1 | 1 | 0.01 | 0 | 0.5 | 0 | 0 | 0.01 | 0.5 |
| WP | 0.01 | 0.5 | 0.01 | 1 | 0.5 | 1 | 0 | 0.01 | 0 | 0 |
| EP | 1 | 0.01 | 0 | 0.5 | 1 | 0 | 0 | 0.01 | 1 | 0 |
| AF | 0 | 0.01 | 0.5 | 1 | 0 | 1 | 1 | 0.5 | 0.01 | 0.01 |
| MD | 0 | 0 | 0 | 0 | 0 | 1 | 1 | 0.5 | 0.01 | 0.01 |
| IN | 0 | 0 | 0 | 0.01 | 0.01 | 0.5 | 0.5 | 1 | 0.1 | 0.01 |
| SEA | 0.01 | 0 | 0.01 | 0 | 1 | 0.01 | 0.01 | 0.1 | 1 | 0.01 |
| AU | 0.01 | 0 | 0.5 | 0 | 0 | 0.01 | 0.01 | 0.01 | 0.01 | 1 |

| **80 - 100 Myr** | WN | EN | NT | WP | EP | AF | MD | IN | SEA | AU |
|---|---|---|---|---|---|---|---|---|---|---|
| WN | 1 | 0.5 | 0.01 | 0 | 0.5 | 0 | 0 | 0 | 0.01 | 0 |
| EN | 0.5 | 1 | 0.01 | 1 | 0.1 | 0.01 | 0 | 0 | 0 | 0 |
| NT | 0.01 | 0.01 | 1 | 0.01 | 0 | 0.5 | 0 | 0 | 0 | 0.5 |
| WP | 0 | 1 | 0.01 | 1 | 0.5 | 0.5 | 0 | 0 | 0 | 0 |
| EP | 0.5 | 0.1 | 0 | 0.5 | 1 | 0 | 0 | 0 | 1 | 0 |
| AF | 0 | 0.01 | 0.5 | 0.5 | 0 | 1 | 1 | 1 | 0.01 | 0.01 |
| MD | 0 | 0 | 0 | 0 | 0 | 1 | 1 | 1 | 0 | 0.01 |
| IN | 0 | 0 | 0 | 0 | 0 | 1 | 1 | 1 | 0 | 0.5 |
| SEA | 0.01 | 0 | 0 | 0 | 1 | 0.01 | 0 | 0 | 1 | 0.01 |
| AU | 0 | 0 | 0.5 | 0 | 0 | 0.01 | 0.01 | 0.5 | 0.01 | 1 |

**Table SIV. 2.** Adjacency matrices for five time-intervals used for ancestral biogeographic state estimation using DECX. WN=Western Nearctic, EN=Eastern Nearctic, NT=Neotropic, WP=Western Palearctic, EN=Eastern Palearctic, AF=Afrotropic, MD=Madagascar, IN=India, SEA=South East Asia, AU=Australasia.

| **0 - 10 Myr** | WN | EN | NT | WP | EP | AF | MD | IN | SEA | AU |
|---|---|---|---|---|---|---|---|---|---|---|
| WN | 1 | 1 | 1 | 0 | 1 | 0 | 0 | 0 | 0 | 0 |
| EN | 1 | 1 | 1 | 1 | 0 | 0 | 0 | 0 | 0 | 0 |
| NT | 1 | 1 | 1 | 0 | 0 | 0 | 0 | 0 | 0 | 0 |
| WP | 0 | 1 | 0 | 1 | 1 | 1 | 0 | 1 | 0 | 0 |

| | | | | | | | | | | |
|---|---|---|---|---|---|---|---|---|---|---|
| EP | 1 | 0 | 0 | 1 | 1 | 0 | 0 | 1 | 1 | 0 |
| AF | 0 | 0 | 0 | 1 | 0 | 1 | 1 | 1 | 0 | 0 |
| MD | 0 | 0 | 0 | 0 | 0 | 1 | 1 | 0 | 0 | 0 |
| IN | 0 | 0 | 0 | 1 | 1 | 1 | 0 | 1 | 1 | 0 |
| SEA | 0 | 0 | 0 | 0 | 1 | 0 | 0 | 1 | 1 | 1 |
| AU | 0 | 0 | 0 | 0 | 0 | 0 | 0 | 0 | 1 | 1 |

| **10 - 30 Myr** | WN | EN | NT | WP | EP | AF | MD | IN | SEA | AU |
|---|---|---|---|---|---|---|---|---|---|---|
| WN | 1 | 1 | 1 | 0 | 1 | 0 | 0 | 0 | 0 | 0 |
| EN | 1 | 1 | 1 | 1 | 0 | 0 | 0 | 0 | 0 | 0 |
| NT | 1 | 1 | 1 | 0 | 0 | 1 | 0 | 0 | 0 | 0 |
| WP | 0 | 1 | 0 | 1 | 1 | 1 | 0 | 1 | 0 | 0 |
| EP | 1 | 0 | 0 | 1 | 1 | 0 | 0 | 1 | 1 | 0 |
| AF | 0 | 0 | 1 | 1 | 0 | 1 | 1 | 1 | 0 | 0 |
| MD | 0 | 0 | 0 | 0 | 0 | 1 | 1 | 0 | 0 | 0 |
| IN | 0 | 0 | 0 | 1 | 1 | 1 | 0 | 1 | 1 | 0 |
| SEA | 0 | 0 | 0 | 0 | 1 | 0 | 0 | 1 | 1 | 1 |
| AU | 0 | 0 | 0 | 0 | 0 | 0 | 0 | 0 | 1 | 1 |

| **30 - 60 Myr** | WN | EN | NT | WP | EP | AF | MD | IN | SEA | AU |
|---|---|---|---|---|---|---|---|---|---|---|
| WN | 1 | 1 | 1 | 0 | 1 | 0 | 0 | 0 | 0 | 0 |
| EN | 1 | 1 | 1 | 1 | 0 | 0 | 0 | 0 | 0 | 0 |
| NT | 1 | 1 | 1 | 0 | 0 | 1 | 0 | 0 | 0 | 1 |
| WP | 0 | 1 | 0 | 1 | 1 | 1 | 0 | 0 | 0 | 0 |
| EP | 1 | 0 | 0 | 1 | 1 | 0 | 0 | 0 | 1 | 0 |
| AF | 0 | 0 | 1 | 1 | 0 | 1 | 1 | 1 | 1 | 0 |
| MD | 0 | 0 | 0 | 0 | 0 | 1 | 1 | 1 | 0 | 0 |
| IN | 0 | 0 | 0 | 0 | 0 | 1 | 1 | 1 | 1 | 0 |
| SEA | 0 | 0 | 0 | 0 | 1 | 1 | 0 | 1 | 1 | 1 |
| AU | 0 | 0 | 1 | 0 | 0 | 0 | 0 | 0 | 1 | 1 |

| **60 - 80 Myr** | WN | EN | NT | WP | EP | AF | MD | IN | SEA | AU |
|---|---|---|---|---|---|---|---|---|---|---|
| WN | 1 | 1 | 0 | 0 | 1 | 0 | 0 | 0 | 0 | 0 |
| EN | 1 | 1 | 0 | 1 | 0 | 0 | 0 | 0 | 0 | 0 |
| NT | 0 | 0 | 1 | 0 | 0 | 1 | 0 | 0 | 0 | 1 |
| WP | 0 | 1 | 0 | 1 | 1 | 1 | 0 | 0 | 0 | 0 |
| EP | 1 | 0 | 0 | 1 | 1 | 0 | 0 | 0 | 1 | 0 |
| AF | 0 | 0 | 1 | 1 | 0 | 1 | 1 | 1 | 0 | 0 |
| MD | 0 | 0 | 0 | 0 | 0 | 1 | 1 | 1 | 0 | 0 |
| IN | 0 | 0 | 0 | 0 | 0 | 1 | 1 | 1 | 0 | 0 |
| SEA | 0 | 0 | 0 | 0 | 1 | 0 | 0 | 0 | 1 | 0 |
| AU | 0 | 0 | 1 | 0 | 0 | 0 | 0 | 0 | 0 | 1 |

| **80 - 100 Myr** | WN | EN | NT | WP | EP | AF | MD | IN | SEA | AU |
|---|---|---|---|---|---|---|---|---|---|---|
| WN | 1 | 1 | 0 | 0 | 1 | 0 | 0 | 0 | 0 | 0 |
| EN | 1 | 1 | 0 | 1 | 1 | 0 | 0 | 0 | 0 | 0 |
| NT | 0 | 0 | 1 | 0 | 0 | 1 | 0 | 0 | 0 | 1 |
| WP | 0 | 1 | 0 | 1 | 1 | 1 | 0 | 0 | 0 | 0 |
| EP | 1 | 1 | 0 | 1 | 1 | 0 | 0 | 0 | 1 | 0 |
| AF | 0 | 0 | 1 | 1 | 0 | 1 | 1 | 1 | 0 | 1 |
| MD | 0 | 0 | 0 | 0 | 0 | 1 | 1 | 1 | 0 | 1 |
| IN | 0 | 0 | 0 | 0 | 0 | 1 | 1 | 1 | 0 | 1 |
| SEA | 0 | 0 | 0 | 0 | 1 | 0 | 0 | 0 | 1 | 0 |

| | | | | | | | | | | |
|---|---|---|---|---|---|---|---|---|---|---|
| AU | 0 | 0 | 1 | 0 | 0 | 1 | 1 | 1 | 0 | 1 |

**Figure SIV. 1**. (see external figures) Historical biogeography estimated by DECX. Pie-charts at the nodes are marginal probabilities of each biogeographic area, i.e., the relative frequency of an area summed across all probable ranges. WN=Western Nearctic, EN=Eastern Nearctic, NT=Neotropic, WP=Wetern Palearctic, EN=Eastern Palearctic, AF=Afrotropic, MD=Madagascar, IN=India, SEA=South East Asia, AU=Australasia.

### V - Biogeographic patterns of diversification: combining BAMM and DECX

We investigated the contribution of the three main processes explaining a regional species pool (dispersal, speciation, and extinction) to understand the disparity in extant species richness among biogeographic regions. We combined the speciation and extinction rates estimates for ancestral lineage obtained from BAMM with the biogeographic ranges and timing of dispersal events estimated with DECX to compare the accumulation of “out-of” *versus* “into” dispersal events, the average speciation rates and the average extinction rates among regions. We also integrated time in this analysis by computing these statistics for four different time intervals.

#### *1 - Dispersal events*

We divided time into four intervals: from 84 to 56 Myrs (Cretaceous-Paleocene: 28 Myrs-long), from 56 to 33.9 Myrs (Eocene: ~22 Myrs-long), from 33.9 to 23 Myrs (Oligocene: ~11 Myrs-long), and from 23 to 0 Myrs (Miocene-Present: ~23 Myrs-long). For each period of time we computed the sum of dispersal events among biogeographic regions. DECX does not implement biogeographic stochastic mapping and does not allow a simple use of ancestral state uncertainty in further analyses. Hence, we extracted the ranges with the highest probability at the nodes and identified the branches along which a dispersal event (range expansion) occurred. Then, to obtain a timing of dispersal events, we drew 1000 random timing of events along the branches for each event. For each replicate, we recorded the number of dispersal events between biogeographic regions occurring during each of the four periods of time. When a dispersal event occurred from a node occupying multiple areas, the source of the dispersal event was chosen by taking among the source area the one with the highest dispersal multiplier during that time period. In case there was two source areas with an equal multiplier we chose one randomly. Finally, we computed the average number of dispersal events across the 1000 stochastic timing of events and transform these sums of events into percentages of the total number of events during a period of time. For this analysis we reduced the number of areas from nine to six by combining Eastern and Western Nearctic into Nearctic, Eastern and Western Palearctic into Palearctic, India and Southeast Asia into Southeast Asia (Australasia was kept as a single area). Combining these areas improved the visualization of the results without actually loosing significant biogeographic information.

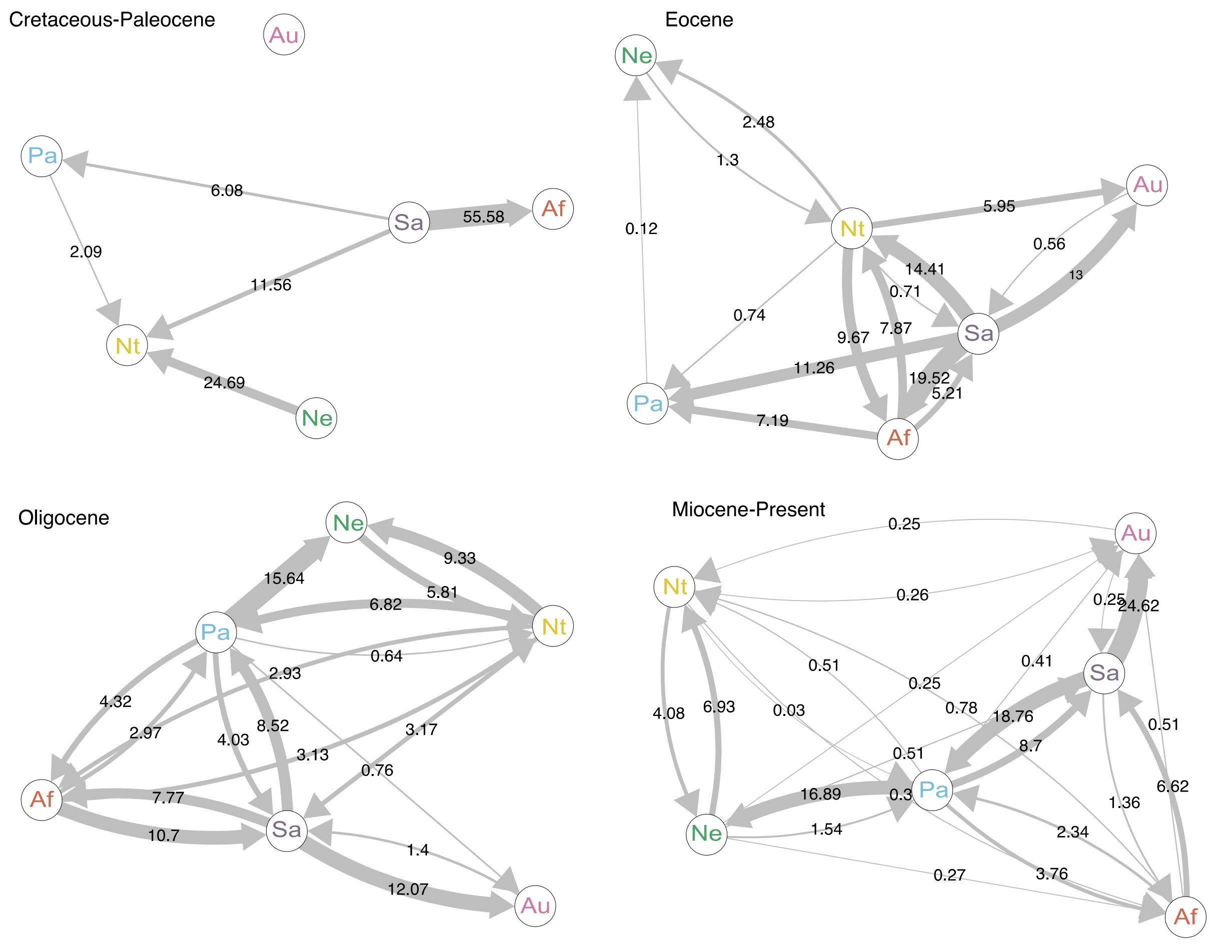


**Figure SV. 1.** Pattern of dispersal events during four time-intervals (Cretaceous-Paleocene, Eocene, Oligocene, Miocene-Present). Arrows indicate the direction of dispersal events. Thickness of the arrow is proportional to the amount of dispersal events occurring during the period of time. The number of dispersal events is the average number over 1000 stochastic timings of events. Numbers indicated are percentages of the total number of dispersal events during a specific period of time.

#### *2 - Lineage-area frequency through time*

We used DECX results to estimate the frequency of lineages sampled in the tree in each area through time. The number of lineages was computed within 0.5 My time-intervals. Here, in case of a dispersal event occurring along a branch, we took the midpoint of the branch as the timing of the dispersal event.

#### *3 - Biogeography and diversification rates*

We estimated how speciation and extinction rates varied through time within each biogeographic region by combing DECX ancestral range estimates and speciation/extinction rates estimation from BAMM.

a) Sliding window analysis

For the speciation and extinction rates we extracted the rates through time for all branches using the function dtRates (BAMMtools, Rabosky et al. 2014). The function dtRates discretized the phylogeny into time bins. Speciation and extinction rates of branches are then extracted for within each time window. The width of time bins is controlled by the parameter tau. In our case, we used a tau of 0.05, i.e., the topology was divided into 20 windows of equal size. In parallel, we inferred the ranges occupied along the branches from our ancestral range estimation (DECX analyses). For each branch we compared the ranges occupied at the upper node to the lower node. In the case of dispersal events occurring along the branch, we sampled another 100 stochastic timings of event for each dispersal event. Before the dispersal event, the lineage occupied the ancestral node ranges, and after the dispersal event, it occupied the descendent node range.

We used a sliding window analysis to obtain diversification rates through time for each biogeographic region. We computed the average speciation, extinction and net diversification rates per region within 4 My time-windows and shifting the window of 1 My. Within a time-window, if a lineage occupied an area, the rates estimated for this branch (or fraction of the branch in case of dispersal events) contributed to the average rate of the region. The average was computed by estimating the number of events in one area (rate*branch length) divided by the sum of branch lengths occupying this area during the same time-interval. The analysis was repeated for 100 stochastic timings of dispersal events.

b) Synthesizing diversification and dispersal

We computed the total number of “into” and “out-of” dispersal events for each biogeographic region within each of the four time-periods (Cretaceous-Paleocene, Eocene, Oligocene, Miocene-Present). Then, we summed for each region across the four time-periods the speciation rate, the extinction rate, the dispersal events into the region and the dispersal events out-of the region. We used this to generate a synthetic visualization of the processes having affected each region since the origin of Nymphalidae (Figure 3).

### VI – Visualization of dispersal and diversification “in real time”

It is difficult to visualize spatial and temporal patterns of diversification on large phylogenetic trees. Ancestral state biogeographic estimation in particular are difficult to synthesize, especially when uncertainty in ancestral ranges is accounted for. Here, we displayed a single realization of historical dispersal events through time on a map with present-day positions of continents for simplicity of implementation. We inferred migrations whenever descendant branches of a node were in different biogeographic regions than its ancestor and used time-stratified matrices to determine which region was more likely to be the source in cases where the ancestral branch was present in more than one location. When there were two or more regions that are equally likely to be the source according to the migration preference matrix we picked one at random.

Lines indicating migrations depart their region of origin at the time when the migrating branch begins (i.e., when the parental node splits into two). The tail of the migration line departs its origin when the head of the lineage is 30% of the way to its destination and all migrations are shown as taking the same amount of time. Regions without any lineages are displayed as white until colonization occurs.

The script used to generate the video is adapted from prior work (Dudas et al. 2017), which uses the baltic module (https://github.com/evogytis/baltic) implemented in python and using matplotlib (cite https://ieeexplore.ieee.org/document/4160265).

Code available here: XXXXXX.

Animation can be seen here: XXXXXX.

We complemented this history of dispersal through time by also displaying the average net diversification rate through time by region estimated in a sliding window analysis (bottom left) and the relative frequency of lineages (sampled in the tree) in each region through time (bottom right).

In the case of net diversification rate, we reduced the number of regions by combining eastern and western Nearctic into "Nearctic", eastern and western Palearctic into "Palearctic", India and South-East Asia into "South-East Asia" and Afrotropics and Madagascar into "Afrotropics". We kept eastern Palearctic, western Palearctic, Afrotropics and Madagascar separated for the lineage-frequency-through-time plot to better visualize their respective contribution to global diversity.

We complemented this history of dispersal through time by also displaying the average net diversification rate through time by region estimated by the sliding window analysis and the relative frequency of lineages (sampled in the tree) in each region through time. In the case net diversification rate, we reduced the number of regions by combining eastern and western Nearctic into “Nearctic”, eastern and western Palearctic into “Palearctic”, India and South-East Asia into “South-East Asia” and Afrotropics and Madagascar into “Afrotropics”. We kept eastern Palearctic, western Palearctic, Afrotropics and Madagascar separated for the *lineage-frequency-through-time* plot to better visualize their respective contribution of the global diversity.

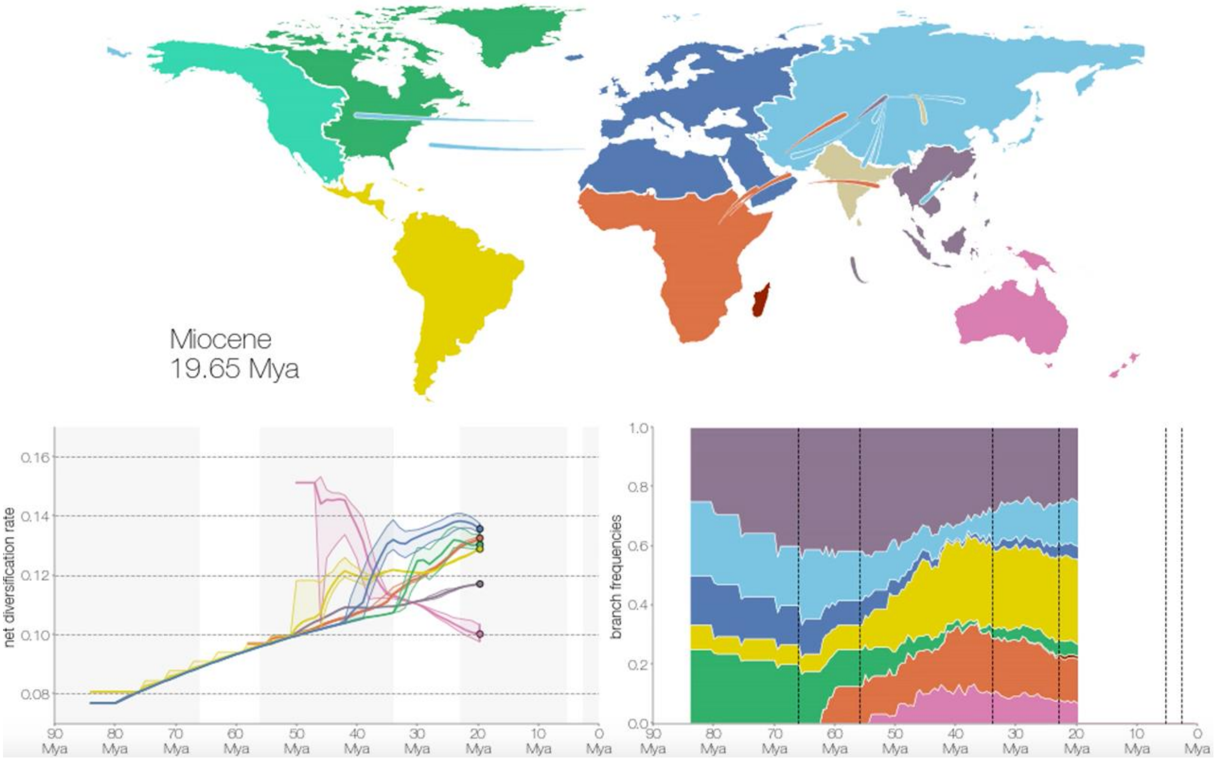


**Figure SVI. 1.** Snapshot of the animation. Upper plot: map of biogeographic regions used in the DECX model. Dispersal events are depicted by the “arrows” travelling from one area to another. Bottom left plot: Net diversification rate through time for the main biogeographic regions, estimated by the sliding window analysis. Bottom right plot: relative frequency lineages sampled in the tree in each biogeographic region through time.

### VII - Biogeographic patterns of diversification: regional diversification

We investigated the biogeographic pattern of diversification using an alternative approach consisting of focusing on the clades having diversified in one region only. We call these events hereafter “regional diversification”.

#### *1 - Dataset*

Using the distribution of extant species and our inferred ancestral ranges we identified these regional diversification events and we computed a set of statistics to understand the processes explaining the current distribution of diversity.

We arbitrarily defined a local diversification event as a clade of at least four tips (i.e., extant taxa included in our tree) that diversified almost entirely in a single biogeographic region. We allowed rare events of range expansion, or recent dispersal events in one additional area. We also had to use a reduced number of regions: Neotropics, Afrotropics, Southeast Asia combined with Australasia, Palearctic, Nearctic. Australasia needed to be combined with South-East Asia because interchanges between the two regions happened too frequently. We identified 90 regional diversification events. For each of these events we estimated the fraction of diversity included in our tree and therefore the total number of extant species. This set of local diversifications accounted for an estimated 5373 species (and 2623 included tips) and ca. 90% of the estimated total diversity (92% of tips included in the tree).

#### *2 - Data analyses*

We designed the following tests to investigate whether (1) any pattern of diversification emerges among regional diversification events belonging to the same biogeographic region, for example in terms of time-variation of speciation rate and whether (2) the age of the clade, recent dynamics of diversification or dynamics occurring during the early stage of diversification explain the extant diversity of these radiations.

We fitted a model for each diversification event independently, in which speciation rate was modelled as an exponential function of time, while extinction rate remained constant (Morlon et al. 2011) using the R-package RPANDA 1.5 (Morlon et al. 2016). We specified the appropriate sampling fraction and excluded the stem branch in the estimation of parameters. For each clade we estimated the extinction parameter ($\mu$) and the two parameters for speciation: speciation rate at present ($\lambda_0$) and coefficient of time variation ($\alpha$). Using these parameters, we also computed the speciation rate at the crown age ($\lambda_{crown}$), the net diversification at present ($netDiv_0$) and the net diversification rate at the crown age ($netDiv_{crown}$). We tested for a relationship between crown age, $netDiv_0$, $netDiv_{crown}$ or $\alpha$ and the number of extant species (log-transformed species richness) by fitting a linear model for each biogeographic region.

Crown age had a strong significant effect on the number of extant species. Therefore, we ran the linear models again for each region on the residual species richness after removing the effect of crown age. We also removed the Neotropical radiation of *Vanessa* butterflies of only five species, which was characterized by an extremely high $netDiv_{crown}$. We show both tests, with and without this clade.

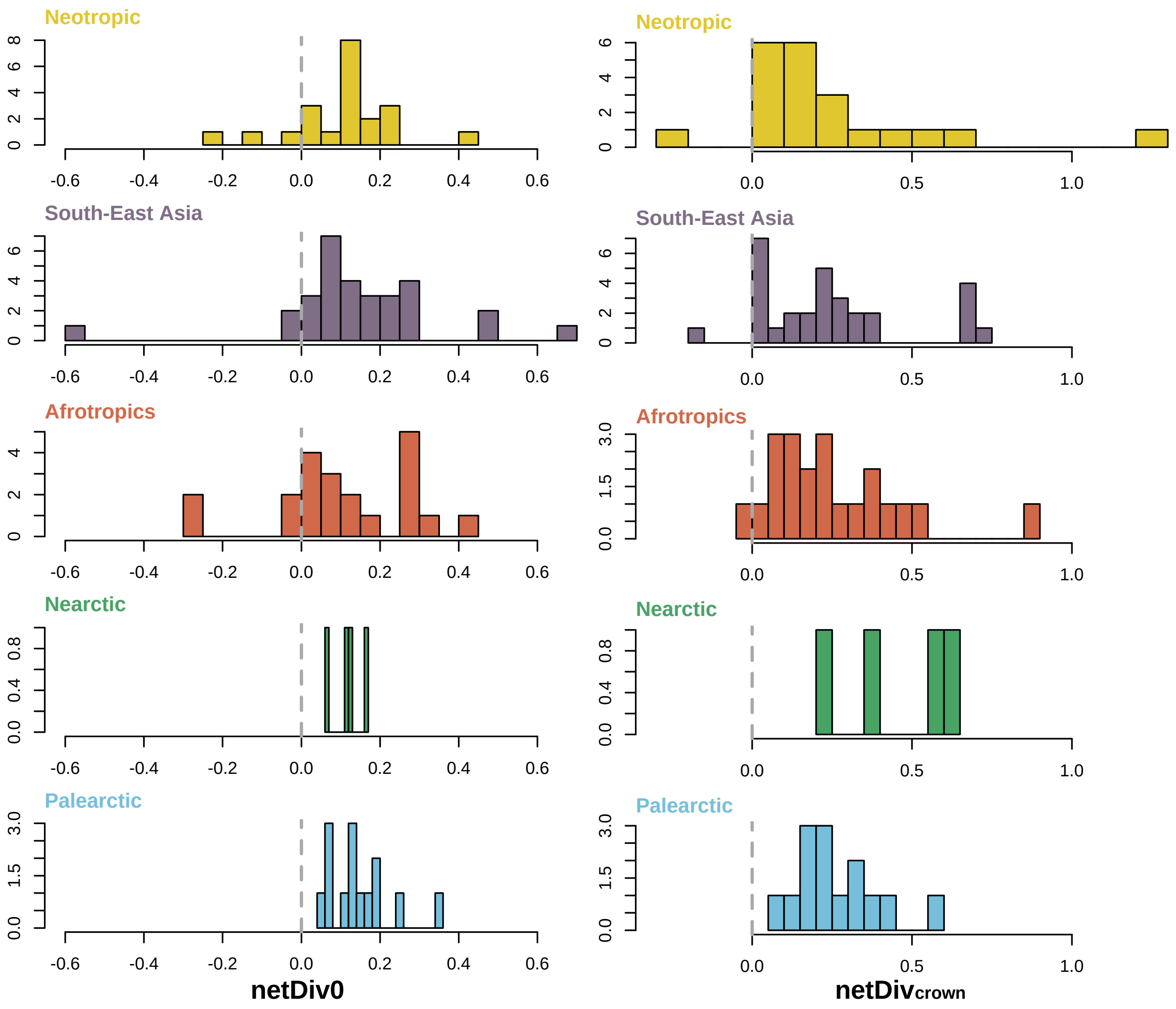


**Figure SVII. 1.** Distribution of net diversification rate at present (netDIv$_0$) and net diversification at the crown ($\lambda_{crown}$) for the regional diversification events.

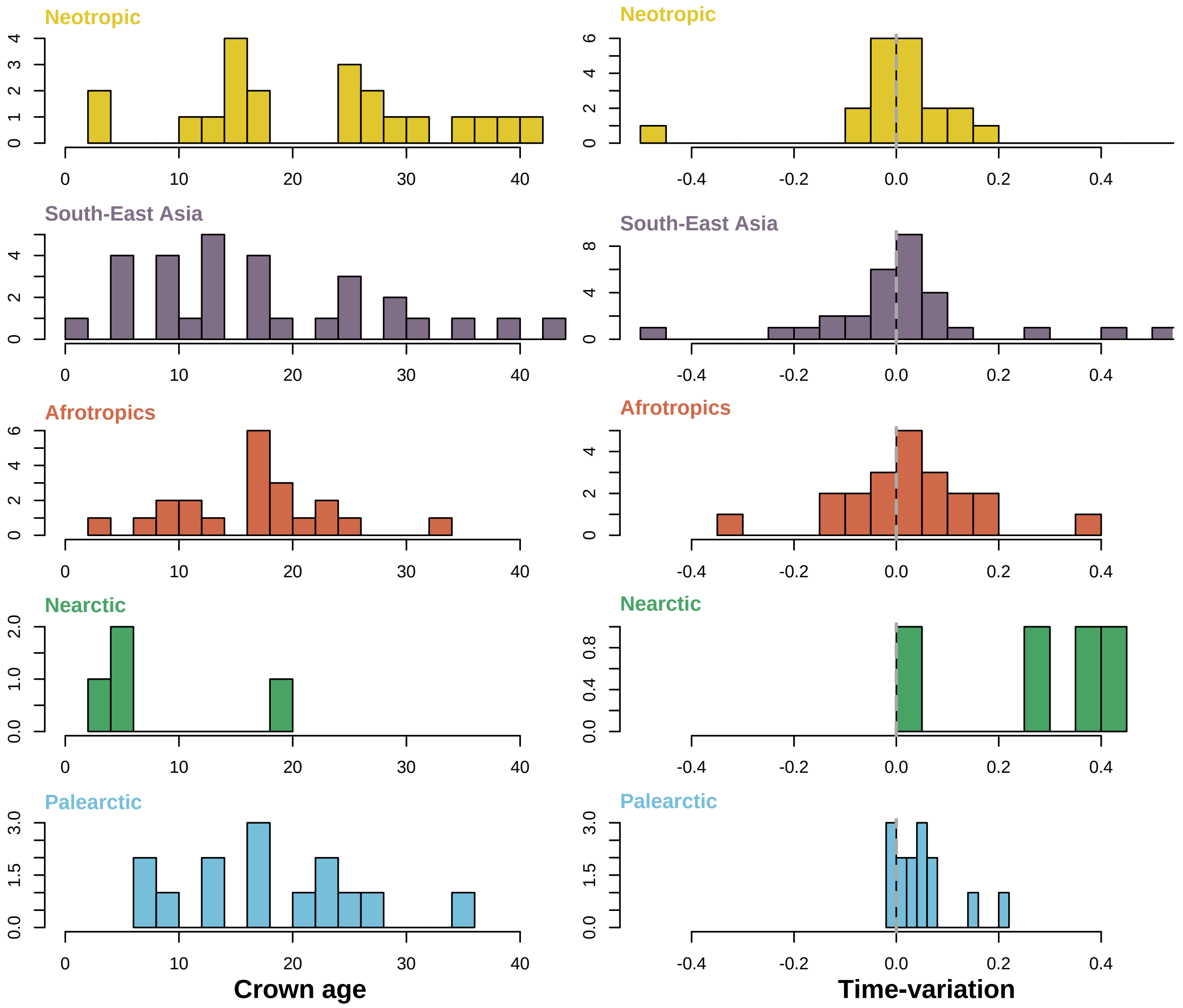


**Figure SVII. 2.** Distribution of crown age and time-variation parameter (α) for the regional diversification events. A positive α indicates a rate decreasing through time, a negative α indicates a rate increasing through time.

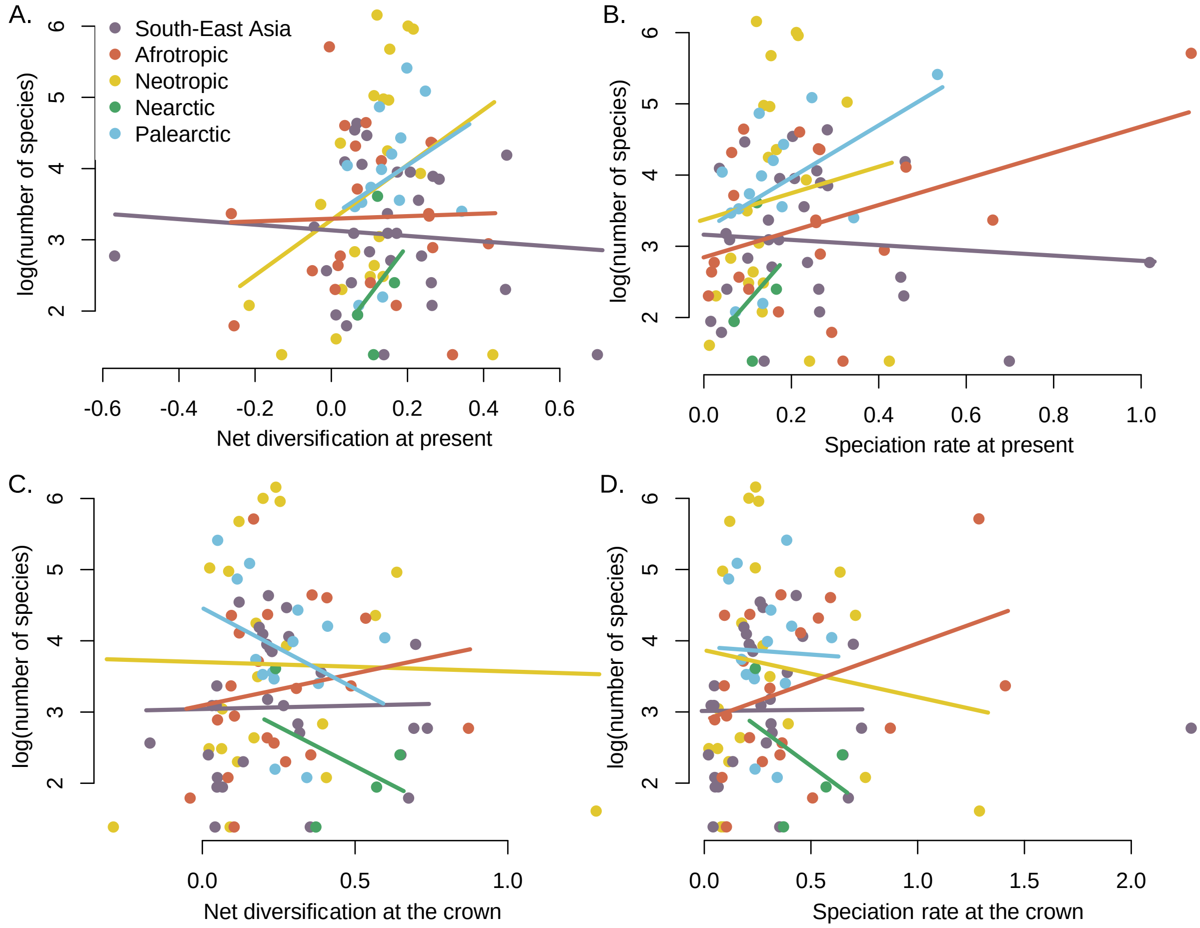


**Figure SVII. 3.** A. Number of species as a function of net diversification rate at present (netDIv$_0$). B. Number of species as a function of speciation rate at present ($\lambda_0$). C. Number of species as a function of net diversification rate at the crown (netDIv$_{crown}$). D. Number of species as a function of speciation rate at present ($\lambda_{crown}$).

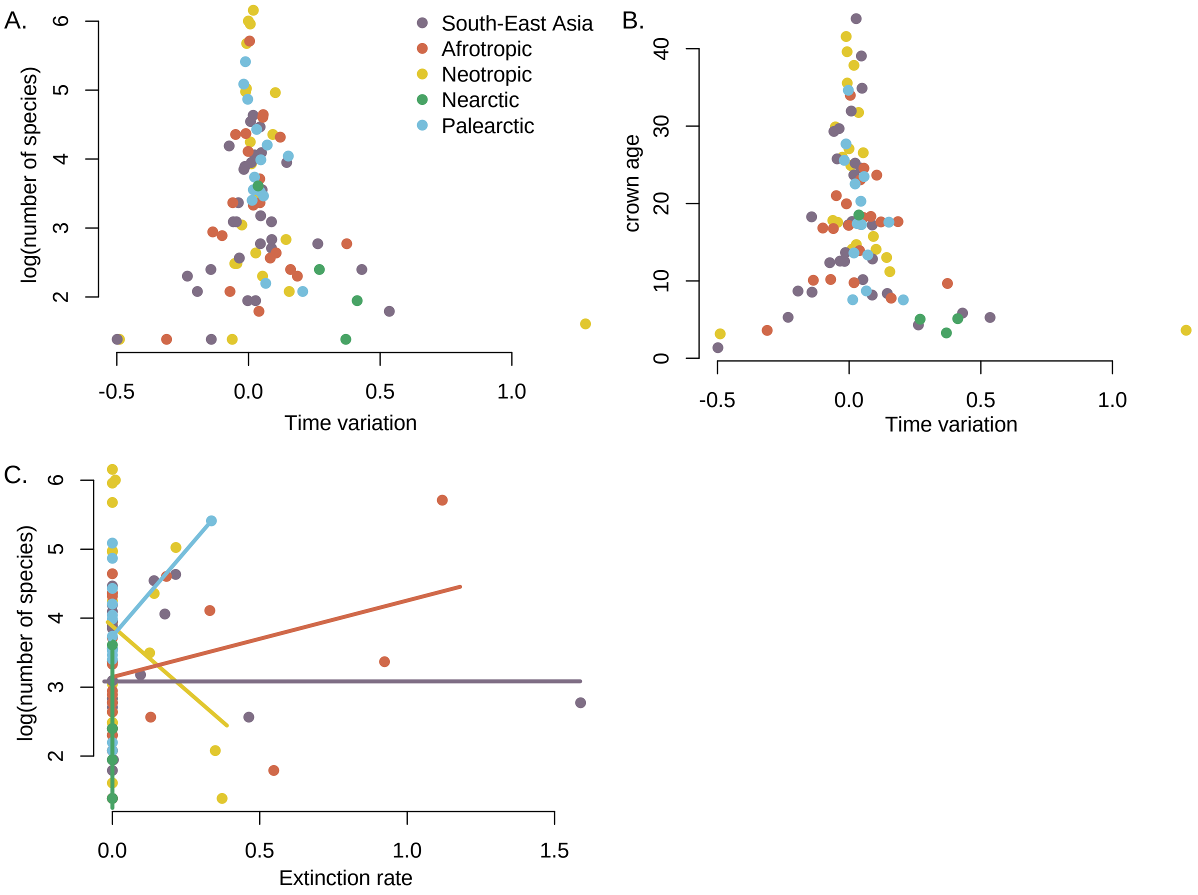


**Figure SVII. 3.** A. Number of species as a function of the time-variation parameter (α). A positive α indicates a rate decreasing through time, a negative α indicates a rate increasing through time. B. Crown age as a function of the time-variation parameter (α). C. Number of species as a function of extinction rate (μ).

**Table SVII. 1.** Results of the of linear model fitted between species richness (on log scale) and crown age, $netDiv_0$ or $netDiv_{crown}$,

| | | Estimate | Std.Error | t-value | p-value |
|---|---|---|---|---|---|
| Neotropics | Intercept | 0.04506 | 0.73845 | 0.061 | 0.952057 |
| | crown age | 0.11916 | 0.02398 | 4.969 | **0.000117** |
| | netdiv_root | 2.14988 | 0.86593 | 2.483 | **0.023772** |
| | netdiv_pres | 4.27479 | 1.79134 | 2.386 | **0.028912** |
| South-East Asia | Intercept | 1.71019 | 0.61326 | 2.789 | 0.00977 |
| | crown age | 0.05121 | 0.01971 | 2.598 | **0.01525** |
| | netdiv_root | 1.28347 | 0.86358 | 1.486 | 0.14925 |
| | netdiv_pres | 0.9995 | 0.96884 | 1.032 | 0.31174 |
| Afrotropics | Intercept | 0.17648 | 0.50327 | 0.351 | 0.7301 |
| | crown age | 0.14328 | 0.02189 | 6.545 | **5.00E-06** |
| | netdiv_root | 1.94088 | 0.72367 | 2.682 | **0.0158** |
| | netdiv_pres | 2.53985 | 0.89855 | 2.827 | **0.0116** |
| Palearctic | Intercept | -0.18282 | 0.72376 | -0.253 | 0.8057 |
| | crown age | 0.12955 | 0.01962 | 6.602 | **6.07E-05** |
| | netdiv_root | 2.79535 | 1.13264 | 2.468 | **0.03322** |
| | netdiv_pres | 6.2119 | 1.58021 | 3.931 | **0.00282** |
| Nearctic | Intercept | 1.33119 | 0.32695 | 4.072 | 0.0554 |
| | crown age | 0.12562 | 0.03247 | 3.869 | 0.0608 |

**Table SVII. 2.** Results of the of linear model fitted between the residual species richness (after removing the effect of time) and $netDiv_0$ and $netDiv_{crown}$,

| | | Estimate | Std.Error | t-value | p-value |
|---|---|---|---|---|---|
| Neotropics | Intercept | -0.8229 | 0.3588 | -2.293 | 0.0341 |
| | netDivcrown | 1.6524 | 0.7644 | 2.162 | **0.0443** |
| | netdiv_pres | 4.1827 | 1.8089 | 2.312 | **0.0328** |
| South-East Asia | Intercept | -0.2744 | 0.2829 | -0.97 | 0.341 |
| | netDivcrown | 0.8143 | 0.717 | 1.136 | 0.266 |
| | netdiv_pres | 0.4789 | 0.8083 | 0.592 | 0.558 |
| Afrotropics | Intercept | -0.6722 | 0.2596 | -2.589 | 0.0185 |
| | netDivcrown | 1.7866 | 0.7148 | 2.499 | **0.0223** |
| | netdiv_pres | 2.2009 | 0.8506 | 2.587 | **0.0186** |
| Palearctic | Intercept | -1.1453 | 0.4085 | -2.803 | 0.01717 |
| | netDivcrown | 1.3908 | 0.9761 | 1.425 | 0.18195 |
| | netdiv_pres | 5.2636 | 1.6821 | 3.129 | **0.00959** |
| Nearctic | Intercept | -0.9797 | 0.6292 | -1.557 | 0.363 |
| | netDivcrown | 1.0095 | 0.9311 | 1.084 | 0.474 |
| | netdiv_pres | 4.4365 | 4.3557 | 1.019 | 0.494 |
