## Supplementary figures and images for "The latitudinal diversity gradient in brush-footed butterflies (Nymphalidae): conserved ancestral tropical niche but different continental histories"

### Figure SIII. 1

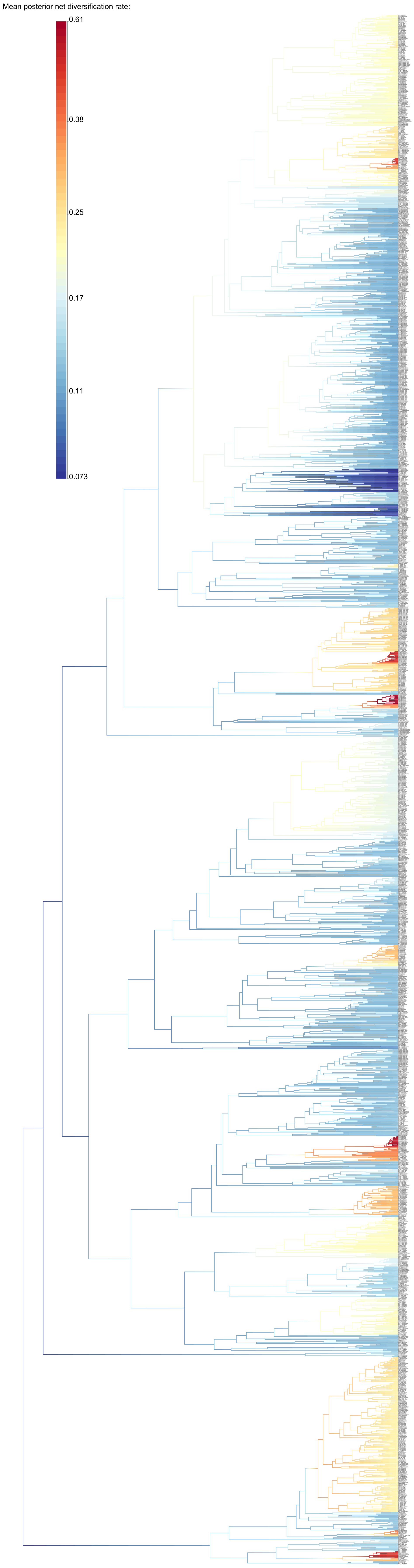

### Figure SIV. 1

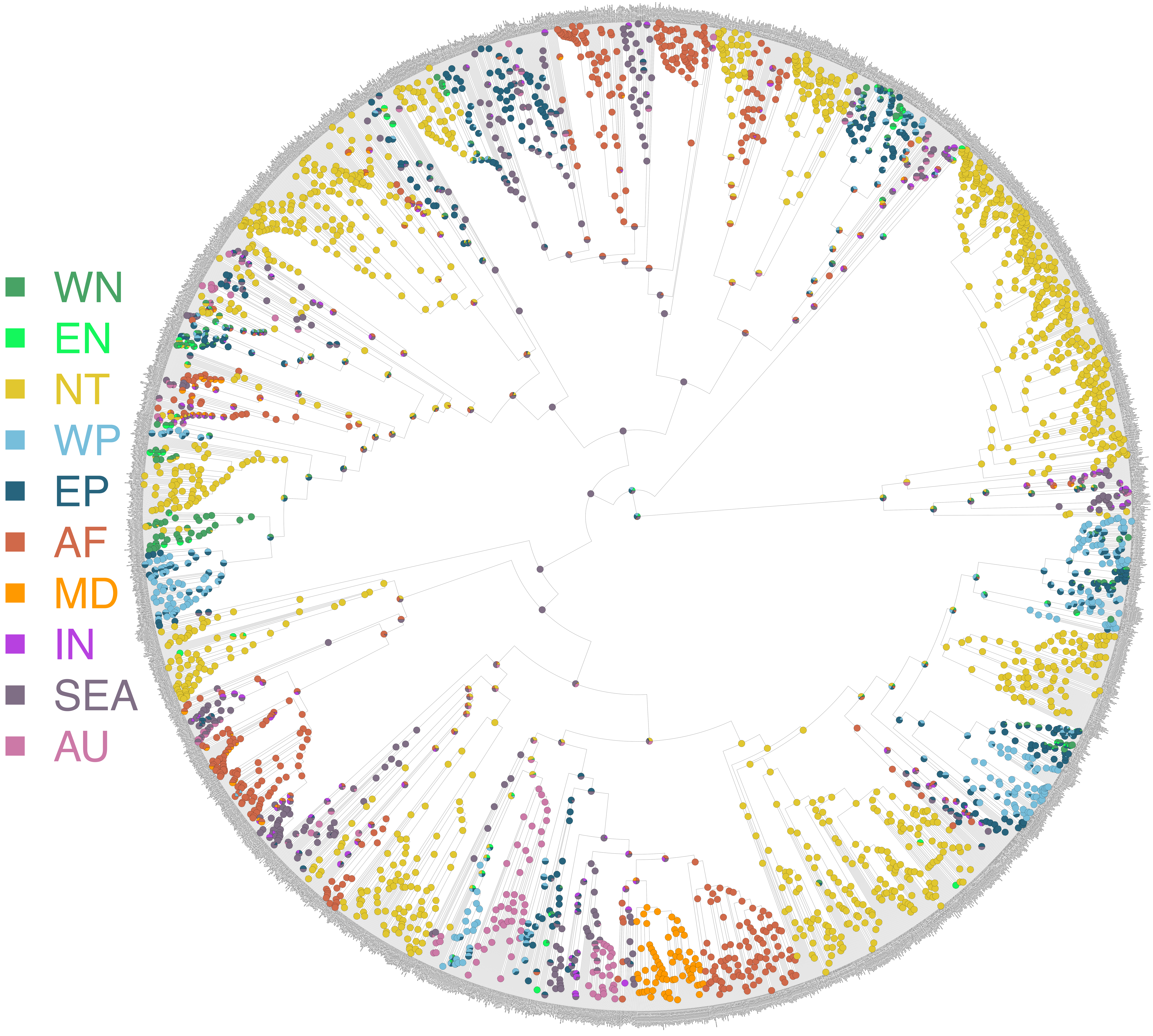
